# Carotid body mitochondria exhibit normal oxygen affinity despite COX4I2 enrichment

**DOI:** 10.64898/2026.06.26.734739

**Authors:** Agnieszka Swiderska, Michael P. Murphy, Gina L. J. Galli, Andrew W. Trafford

**Affiliations:** Unit of Cardiac Physiology, Division of Cardiovascular Sciences, School of Medical Sciences, Faculty of Biology, Medicine and Health, University of Manchester, Core Technology Facility, Manchester, United Kingdom; Medical Research Council Mitochondrial Biology Unit and Department of Medicine, University of Cambridge, Cambridge, United Kingdom

**Keywords:** Carotid body, oxygen sensing, left ventricle, mitochondria, mitochondrial reactive oxygen species, heart disease

## Abstract

The carotid body (CB) is the key peripheral oxygen sensor. CB mitochondria are hypothesised to be uniquely adapted with unusually low intrinsic oxygen affinity which, in association with nitric oxide (NO) and reactive oxygen species signalling, enables acute responsiveness to hypoxia. However, CB mitochondrial physiology or intrinsic oxygen affinity have never been measured directly. We sought to address this key gap by isolating sheep CB mitochondria and comprehensively characterising their phenotype and contrasting them to a non-oxygen sensing tissue, left ventricular myocardium (LV).

High resolution respirometry, liquid chromatography mass spectrometry, enzymatic assays and in silico modelling were used to characterise mitochondrial content, aerobic capacity, oxygen affinity, complex subunit abundance and activity, H_2_O_2_ production and NO sensitivity in ovine CB and LV. Mitochondrial oxygen affinity (P_50_ = 0.089 mmHg) was lower in the CB than the LV (P_50_ = 0.058 mmHg; p = 0.005). Whilst mitochondrial content was lower in the CB, CB mitochondria had higher respiratory rates and enzymatic activity than LV. H_2_O_2_ production and NO sensitivity were similar in the two tissues.

While intrinsic mitochondrial oxygen affinity is slightly lower in the oxygen sensing CB than in the non-oxygen sensing LV, this difference is small. Hence, any role of mitochondria in CB oxygen sensing is not due to an intrinsic difference in the O_2_ affinity of cytochrome oxidase due to differential expression of its subunits. Instead, this work suggests that differences in O_2_ affinity in vivo are secondary to other factors, perhaps including NO, that alter mitochondrial O_2_ affinity.

## 1. Introduction

The carotid body (CB) is the main peripheral oxygen sensing organ and responds rapidly to changes in blood oxygen levels^1^. There are two main cell types found in the CB: type I cells (the main sensory cells) and type II cells (supportive cells)^2^. Type I cells are sensitive to oxygen and release neurotransmitters upon exposure to hypoxia, increasing carotid sinus nerve firing to the brain^3^. The increased activity leads to sympathetic activation which increases minute ventilation and redirects blood flow to crucial organs^4^.

Despite decades of research, the identity of the CB oxygen sensor remains elusive. While many candidates have been proposed (for review, see^4-6^), the prevailing theory is that mitochondria in CB type I cells are the site of oxygen sensing and that mitochondria signal to the cell membrane to trigger a neurotransmitter release, a hypothesis referred to as the mitochondria-to-membrane hypothesis. This theory is based on a series of studies that suggest mitochondria in the CB are unique from other tissues^4^. In most cells, mitochondria are the primary site of oxygen consumption for oxidative phosphorylation (OXPHOS). Electrons from substrate oxidation are donated to complex I and II of the mitochondrial electron transport chain and subsequently passed to complex III and IV where they bind to the final electron acceptor, oxygen. The energy released from electron transfer is utilised by complexes I, III and IV to pump protons from the mitochondrial matrix to the intermembrane space, resulting in a transmembrane proton electrochemical gradient. Subsequently, protons flow back down their electrochemical gradient through a fifth protein complex (the ATP synthase, or F_1_F_o_-ATPase), which harvests the energy to generate ATP from ADP and P_i_. When oxygen is limited, electron transport is inhibited, the electrochemical gradient dissipates, and ATP production ceases.

In general, mitochondria have an exceptionally high affinity for oxygen with OXPHOS only becoming compromised when cellular oxygen tension falls below 0.5 mmHg^7,8^. However, it has been suggested that this may not be the case in CB mitochondria with several groups demonstrating that mitochondrial membrane potential collapses in intact CB type I cells at levels of oxygen that are approximately 20 times higher (∼40 mmHg) than other cell types (e.g. 2 mmHg in chromaffin cells)^9,10^. Genetic studies revealed that there is a high expression of the mitochondrial complex IV subunit, COX4I2, in CB type I cells^11,12^ and overexpression of COX4I2 in HEK293 systems has been linked to lower oxygen affinity when compared to cells expressing COX4I1 (P_50_ = 0.9 mmHg vs 0.48 mmHg)^13^. This led to the hypothesis that increased COX4I2 is driving low oxygen affinity of type I cell mitochondria. This is pertinent to a putative role in oxygen-sensing because mitochondria bind oxygen within the CuB/haem a3 (cytochrome a3) binuclear centre of complex IV. Importantly, cyanide (complex IV inhibitor) mimics the response to hypoxia in type I cells^12,14^. These studies have led to the assumption that complex IV is the site of oxygen-sensing in CB type I cells, due to its unique molecular phenotype. However, the oxygen affinity of CB mitochondria has only ever been measured indirectly using intact type I cells, so it is possible that other factors could be modulating complex IV activity under hypoxic conditions.

One possible candidate that could be modulating complex IV is nitric oxide (NO), a competitive inhibitor of complex IV and known modulator of electron transport chain activity^15,16^. Previous studies showed that mice lacking neuronal NO synthase (NOS1) were unable to respond to hypoxia^17^, whereas mice lacking endothelial NO synthase (NOS3) became more sensitised to hypoxia compared to the wild-type controls^18^. Another study showed that NO moderately enhances CB hypoxic sensitivity^19^. Therefore, it is possible that complex IV oxygen sensing in CB type I cells is controlled via NO signalling, rather than (or in addition to) an inherently low complex IV oxygen affinity. To shed light on this matter, it is necessary to measure complex IV oxygen affinity in isolated mitochondria, in the absence of any cytosolic signalling.

In addition to the identity of the oxygen sensor, there has been much debate about the mechanism by which the signal is transmitted to the membrane to trigger neurotransmitter release. According to mitochondria-to-membrane hypothesis, reactive oxygen species (ROS) generated by the electron transport chain serve as second messengers in a process known as reverse electron transport (RET) signal to the type I cell membrane to initiate the chemotransduction cascade. Briefly, during hypoxic exposure, there is a partial or full inhibition of complex IV, and flow of electrons ^20^, leading to the accumulation of reduced ubiquinone in CB mitochondria. In the presence of reduced ubiquinone and substrate entry to complex II, electrons flow back to complex I ^20,21^ leading to an increase in both ROS production and NADH ^21^. In CB, increased ROS through RET is proposed to reduce the open probability of potassium channels on the type I cell surface membrane and cause membrane depolarisation, opening of L-type calcium channels and rapid calcium influx. This cascade culminates in neurotransmitter release from type I cells to initiate sympathetic activation and the cardiorespiratory response to hypoxia. Although the mitochondria-to-membrane hypothesis is appealing; there is a surprising lack of studies performed on CB isolated mitochondria given their hypothesised gatekeeper role.

Given the small size of the CB (i.e. rat CB weighs 60 μg^1^), it is necessary to use a large animal model to isolate a sufficient yield of CB mitochondria for downstream analysis. Therefore, the main goals of this study were threefold: i) to develop a protocol for isolating mitochondria from the sheep CB, ii) to characterise their functional phenotype (e.g. aerobic capacity, sites of ROS production and oxygen affinity) and, iii) compare these parameters to mitochondria from a non-oxygen sensing tissue – the left ventricle (LV). To our knowledge, this is the first study to assess CB mitochondrial function directly. Although we find that complex IV oxygen affinity in the CB is lower than the LV, it is within the normal range of mammalian mitochondria and substantially higher than the oxygen affinity of intact type I cells. Thus, intrinsic mitochondrial oxygen affinity alone cannot explain CB oxygen sensing. We also show that isolated CB mitochondria have a higher aerobic capacity than ventricular tissue, but a similar basal H_2_O_2_ production and NO sensitivity.

## 2. Materials and methods

### 2.1. Animal model

All work involving the use of animals was performed in accordance with The Animals (Scientific Procedures) Act 1986 and EU Directive 2010/63. Local ethical approval was obtained from The University of Manchester Animal Welfare and Ethical Review Board. The reporting of use of animals in line with ARRIVE2 guidelines^22^. All procedures were performed using female Welsh Mountain sheep (47 animals, ∼ 18 - 24 months of age). Animals were fed ad libitum hay supplemented with ruminant concentrate and maintained at ∼ 21°C on a 12:12 hour light:dark cycle. No animals were excluded from the study or post data acquisition analysis.

### 2.2. Tissue dissection

Sheep were injected intravenously with 20,000 IU heparin prior to euthanasia with intravenous sodium pentobarbitone (Pentoject; 200 mg.kg^-1^) to prevent coagulation and improve post-mortem tissue viability. Firstly, the heart was rapidly excised, and a myocardial section of LV was taken from the mid-posterior wall. Secondly, an incision was made along the jugular groove to expose the common carotid artery and its bifurcation. The common carotid artery and bifurcation were dissected out. The CBs were then identified and dissected out of the tissue. Samples were taken for mitochondrial isolation, homogenate preparation or snap frozen and stored in liquid nitrogen for subsequent molecular analyses.

### 2.3. Mitochondrial isolation

Mitochondrial isolation was performed using differential centrifugation^23^. Briefly, tissue was minced in ice-cold MSE isolation buffer (containing (in mmol.L^-1^): D-mannitol, 220; sucrose, 70; 3-(N-Morpholino)propanesulfonic acid (MOPS), 5; ethylenediaminetetraacetic acid disodium salt dihydrate (NaEDTA), 2; and 0.1% bovine serum albumin (BSA; pH = 7.35 at 4°C)) and incubated with trypsin (0.75 mg.mL^-1^, T0303, Sigma-Aldrich) for 10 minutes on ice, followed by trypsin inhibitor (0.22 mg.mL^-1^, T9128, Sigma-Aldrich) for another 3 minutes. Tissue was then resuspended in fresh MSE solution and homogenised using a motorised Teflon pestle. The solution was transferred into a falcon tube and centrifuged at 800 x g for 10 minutes at 4°C. The suspension was then filtered through a surgical gauze and centrifuged at 9000 x g for 10 minutes at 4°C. For LV samples, the resulting mitochondrial pellet was resuspended in fresh MSE buffer and centrifuged a second time at 9000 x g for 10 minutes. This step was not possible for CB mitochondria because the pellet was much smaller, and too much tissue was lost on a second spin. After the final spin, the CB and LV pellets were resuspended in fresh MSE buffer until experimentation.

### 2.4. Mitochondrial oxygen consumption and ROS production in isolated mitochondria

Mitochondrial respiration and H_2_O_2_ production were measured with an Oroboros Oxygraph-O2K fitted with 0.5 mL chambers.

#### 2.4.1. SUIT protocol

Mitochondrial respiration rate was characterised in different respiratory states by utilizing a substrate-uncoupler-inhibitor-titration (SUIT) protocol. Amplex® UltraRed (10 µmol.L^-1^), 1 U.mL^-1^ horseradish peroxidase (HRP) and 5 U.mL^-1^ superoxide dismutase (SOD) were added into each chamber to measure H_2_O_2_ production. The H_2_O_2_ signals were calibrated by adding 40 μmol.L^-1^ H_2_O_2_ twice to achieve final concentrations of 0.1 and 0.2 μmol.L^-1^ in each chamber.

Mitochondria were injected to each chamber before complex CI substrates were added (pyruvate (6.25 mmol.L^-1^), malate (2 mmol.L^-1^), and glutamate (10 mmol.L^-1^)) to measure LEAK respiration without adenylates (LEAK_CI_). A saturating concentration of ADP (0.5 mmol.L^-1^) was added to measure respiratory rate OXPHOS with CI substrates (OXPHOS_CI_), and succinate (10 mmol.L^-1^) was then added to measure OXPHOS with CI and CII substrates (OXPHOS_CI+CII_). Maximum electron transfer capacity (ETC_CI+CII_) was measured by adding mitochondrial uncoupler carbonyl cyanide p-trifluoro-methoxyphenyl hydrazone (FCCP). FCCP was added stepwise until no further increases in respiration rate were observed (final concentration of 0.5 - 1 µmol.L^-1^). Subsequently, maximum electron transport capacity with complex II substrates only (ETC_CII_) was measured after adding complex I inhibitor, rotenone (0.15 μmol.L^-1^). Complex III was then inhibited with antimycin A (12.5 μmol.L^-1^) and residual non-mitochondrial respiration rate (ROX) was recorded. Complex IV activity was measured by adding the electron donor N,N,NL,NL-tetramethyl-p-phenylenediamine (TMPD, 0.5 mmol.L^-1^) with ascorbate (2 mmol.L^-1^) to prevent TMPD autooxidation. Finally, sodium azide (50 mmol.L^-1^; complex IV inhibitor) was added to inhibit complex IV and measure background nonmitochondrial oxygen consumption.

Given that BSA was present in the final mitochondrial pellets (which interferes with protein quantification, especially in small samples), we normalised CB and LV respiration rates to citrate synthase activity. The respiratory control ratio (RCR; ratio of OXPHOS_CI_ and LEAK_CI_) and OXPHOS coupling-control ratio (1-(L/P), calculated as (1-(LEAK_CI_/OXPHOS_CI_)) were used to describe the ATP production efficiency in mitochondrial samples.

#### 2.4.2. Mitochondrial oxygen affinity and RET capacity

In order to assess mitochondrial O_2_ affinity, the chamber O_2_ concentration was reduced to ∼ 33 mmHg by blowing nitrogen across the surface. Following addition of isolated mitochondria (20 –30µL CB and 2µL of LV), pyruvate (6.25 mmol.L^-1^), malate (2 mmol.L^-1^), glutamate (10 mmol.L^-1^), ADP (0.5 mmol.L^-1^) and succinate (10 mmol.L^-1^) were added to achieve OXPHOS respiration. Respiration was continuously recorded until oxygen concentration reached zero. The oxygen flux was normalised, and then corrected respiration rate was plotted against the oxygen concentration (mmHg). A nonlinear Michaelis–Menten regression curve was fitted and the P_50_ (50% enzyme saturation) was calculated in GraphPad Prism 8 software (GraphPad Software, Inc. USA).

To measure RET capacity, SOD (5 U.mL^-1^), HRP (1 U.mL^-1^) and Amplex®UltraRed (10 µmol.L^-1^) and isolated mitochondria were added into each chamber to measure baseline H_2_O_2_ production. Succinate (5 mmol.L^-1^) was then added to promote RET via complex I. Calibration was conducted once by adding 40 µmol.L^-1^ H_2_O_2_ solution twice to achieve final concentration of 0.1 and 0.2 µmol.L^-1^ in each chamber.

#### 2.4.3. The effects of nitric oxide inhibition and S-nitrosylation on respiratory rates

Isolated mitochondria, complex I substrates (malate 2 mmol.L^-1^, pyruvate 6.25 mmol.L^-1^, and glutamate 10 mmol.L^-1^) and ADP (0.5 mmol.L^-1^) were added to measure OXPHOS_CI_. Following that, either S-nitrosoglutathione (GSNO, 1 mmol.L^-1^), mitoSNO (mSNO, mitochondria-targeted S-nitrosothiol; 10 µmol.L^-1^) or mito-N-acetylpenicillamine (mNAP, a control compound for mSNO; 10 µmol.L^-1^) were added. Both, GNSO and mSNO compounds act to inhibit mitochondrial respiration by either donating NO as a competitive inhibitor of complex IV (GSNO) or s-nitrosylation (mSNO). Finally, either DL-Dithiothreitol (DTT, 1 mmol.L^-1^) or oxyhaemoglobin (Hb(O_2_)_4_; 2 µL) were added to the chamber. DTT was used to break any s-nitrosylation modifications whereas Hb(O_2_)_4_ was used as an oxygen donor.

### 2.5. Mass Spectrometry

Mitochondrial isolates were centrifugated at 10,000 x g for 10 minutes at 4°C. Supernatant was discarded and mitochondrial pellets resuspended in MSE buffer without BSA. Samples were then centrifuged at 10,000 x g for 10 minutes at 4°C. Mitochondrial pellet was kept at -80°C until samples were processed by the Biological Mass Spectrometry (BioMS) Facility.

#### 2.5.1. Protein Extraction and Lysis

Samples were lysed in S-Trap lysis buffer (5% SDS, 50 mM triethylammonium bicarbonate [TEAB]) and sonicated using a Covaris LE220+ Focused-ultrasonicator. Samples were maintained on ice until processing. Sonication was performed for 10 minutes using the Covaris SonoLab 8.2 software with the following acoustic parameters: peak incident power 500 W, duty factor 50%, 100 cycles per burst, and an average power of 250 W. The instrument’s water bath was maintained at 6 °C throughout.

#### 2.5.2. Reduction and Alkylation

Protein lysates were reduced with DTT to a final concentration of 5 mM. Samples were incubated at 60 °C for 10 min in an Eppendorf Thermomixer C. Following reduction, samples were cooled to room temperature and alkylated with iodoacetamide (IAM) to a final concentration of 15 mM. Alkylation was performed in the dark for 30 min at room temperature. Reactions were quenched by addition of DTT equivalent to that used in the reduction step.

Following reduction and alkylation, protein concentration was determined using the Pierce Rapid Gold BCA assay according to the manufacturer’s instructions. From each quantified lysate, a 20 µg aliquot was taken and made up to a final volume of 50 µL in lysis buffer. These normalised aliquots were subsequently used for digestion in S-Trap microplates.

#### 2.5.3. Protein Digestion Using S-Trap Micro Columns

Protein digests were prepared using S-Trap microplates according to the manufacturer’s protocol with minor modifications. Lysates were acidified by addition of 12% phosphoric acid at a 1:10 (v/v) ratio of acid to lysate, yielding a final acid concentration of approximately 1.2%. Acidified samples were then combined with 350 μL of S-Trap binding buffer (90% methanol, 100 mM TEAB, pH 7.1) and loaded directly onto the S-Trap plate. The plate was centrifuged at 1,000 × g for 2 min to bind proteins.

Bound proteins were washed once with 200 μL of methyl tert-butyl ether (MTBE) to remove methanol-insoluble contaminants, followed by three washes with 200 μL of binding buffer, each centrifuged at 1,000 × g for 2 min.

Trypsin digestion was performed by adding 125 μL of digestion buffer (50mM TEAB) containing 2 μg of TPCK-treated trypsin to each sample containing 20 μg of protein, corresponding to a 1:10 (w/w, enzyme:protein) ratio. Digestion was carried out overnight at 37 °C.

Peptides were eluted sequentially with 80 μL digestion buffer, 200 μL of 0.1% (v/v) formic acid, and 55 μL of 30% acetonitrile with 0.1% formic acid, each followed by centrifugation at 1,000 × g for 2 min.

#### 2.5.4. Peptide Desalting

Peptides were desalted using a Cole-Parmer TELOS® neo™ PRP Fixed-Well Plate containing 10 mg of sorbent. The resin was conditioned with 200 μL of 50% acetonitrile, followed by two equilibration washes with 200 μL of 0.1% formic acid in water.

Samples were applied to the conditioned resin and then centrifuged at 200 × g for 1 min. Each sample was washed twice with 200 μL of 0.1% formic acid.

Peptides were eluted with 2 x 50 μL of 30% acetonitrile containing 0.1% formic acid per step, followed by centrifugation at 200 × g for 1 min. The combined eluates (∼100 μL each) were transferred to LC–MS vials and dried to completeness in a vacuum centrifuge. Dried peptides were stored at 4 °C until reconstitution and analysis.

#### 2.5.5. LC separation

The separation was performed on a Thermo Vanquish Neo UHPLC system configured with buffer A as 0.1% formic acid in water and buffer B as 0.1% formic acid in 80% acetonitrile. A specified injection volume was loaded on to the analytical column (IonOpticks Aurora series Rapid TS, 8 cm x 75 µm ID, 1.7 µm C18) kept at 35 °C at an initial rate of 800 nL.min^-1^ which was then dropped to 200 nL.min^-1^ in 0.25 min to rapidly re-pressurise the column. The separation was also started during this time, with a gradient of 1% B to 5% B over this period. The next step was 5% B to 35% B over 20.25 minutes, 35% B to 100% B over 0.5 minutes before washing for 2 minutes at 100% B and dropping down to 1% B in 0.5 minutes. The complete method time was 30 minutes.

#### 2.5.6. MS source

The analytical column was connected to a Thermo Orbitrap Astral mass spectrometer via a Thermo EasySpray Ion, with a nanospray voltage set at 1600 V and the ion transfer tube temperature set to 275 °C.

#### 2.5.7. MS settings (DIA)

Data was acquired in a data dependent manner using a fixed cycle time of 2 sec, an expected peak width of 15 sec and a default charge state of 2. Full MS data was acquired in positive mode over a scan range of 300 to 1750 Th, with a resolution of 120,000, a normalised AGC target of 300% and an automatic max fill time for a single microscan. Fragmentation data was obtained from signals with a charge state of +2 to +4 and an intensity over 5,000 and they were dynamically excluded from further analysis for a period of 15 sec after a single acquisition within a 10ppm window. Fragmentation spectra were acquired with a resolution of 15,000 with a normalised collision energy of 30%, a normalised AGC target of 300%, first mass of 110 Th and a max fill time of 25 mS for a single microscan. All data was collected in profile mode.

Data was acquired in a data independent manner with an expected peak width of 6 sec and a default charge state of 2. Full MS data was acquired in positive mode over a scan range of 350 to 1750 Th, with a resolution of 240,000, a standard (automatic) normalised AGC target and a maximum injection time of 10 ms for a single microscan. Fragmentation data was obtained according to a table of 150 m/z windows, with 3 Th m/z widths. The DIA window type was set to automatic, and window optimisation was selected. These were acquired in the astral, with a normalised collision energy of 25%. The m/z range was 145-1450. A normalised AGC target of 500% was used with a maximum injection time of 7 ms for a single microscan. All data was collected in centroid mode.

The raw data files from the instrument were processed using Biognosys’s Spectronaut software (version 20.2). The data was processed using a directDIA Analysis against the protein database UniProt Proteome (one protein sequence per gene) for Ovis aries (Taxonomic ID 9940) (version 2025-04; 21, 226 sequences) with the BGS Factory Settings (default) schema. This includes the following parameters:

- the enzyme Trypsin (cleaving at Lysine’s and Arginine’s except where the presence of a C-terminal Proline obstructs cleavage)
- the fixed modifications of Carbamidomethyl (+57.021 Da) to Cysteine’s and variable modifications of Acetylation (+42.011 Da) to protein N-terminus and Oxidation (+15.995 Da) to Methionine’s.

### 2.6. RNA in situ hybridization

ViewRNA™ Tissue Fluorescence Assay (Invitrogen) was performed as per manufacturer’s recommendations. Briefly, formalin-fixed paraffin-embedded tissue samples were sectioned at 5 µm and deparaffinised using Histo-Clear (National Diagnostics, UK). Sections were incubated in Pre-Treatment solution at 90-95°C for 20 minutes, digested by using protease for 15 minutes at 40°C, and fixed again with 10% neutral buffered formalin. Once fixed, target specific ViewRNA probes were applied onto the tissue for 2 hours at 40°C. Two step hybridisation was then performed by incubating the tissue with PreAmplifier and Amplifier Mix, respectively for 30 minutes each at 40°C. ViewRNA probes were labelled with a fluorescent tag (30-minute incubation at 40°C). ReadyProbes™ Tissue Autofluorescence Quenching kit (Invitrogen) was utilised to reduce tissue autofluorescence. Finally, DAPI was used to counterstain cell nuclei and samples were mounted on glass slides using ProLong Glass Antifade Mountant. Slides were then imaged using an Evolve 512 EMCCD camera (Photometrics, UK) on a Zeiss Axiobserver.Z1 inverted microscope (Zeiss, GmbH, Germany) using a Lumencor SpectraX LED light source (395/25, 475/28 and 640/30 nm excitation wavelengths, Lumencor, Orgenon USA).

### 2.7. Spectrophotometric analysis of enzymatic activities

Enzymatic activity of citrate synthase and mitochondrial complexes were measured as previously described^24^ with a BioTek Synergy HTX 301 multimode reader (BioTek, Swindon, UK). Fresh CB and LV tissue were homogenised using a hand-held homogeniser in an ice-cold Mir05 buffer containing (in mmol.L^-1^): sucrose, 110; K-MES, 60; 4-(2-hydroxyethyl)-1-piperazineethanesulfonic acid, 20; monopotassium phosphate, 10; magnesium chloride, 3; ethylene glycol-bis(ß-aminoethyl ether)-N,N,N’,N’-tetraacetic acid, 0.5; and 0.1% BSA. The suspension was then filtered through a 200 µm nylon mesh. Tissue homogenates were kept at -80°C until the day of the assay. Citrate synthase activity was determined in isolated mitochondria or homogenate samples in assay buffer containing: 100 mmol.L^-1^ Tris, 300 µmol.L^-1^ acetyl coenzyme-A, 100 µmol.L^-1^ 5,5L-dithiobis-(2-nitrobenzoic acid)[DTNB] and 0.1% Triton-X-100) by adding oxaloacetate (0.5 mmol.L^-1^). Following oxaloacetate addition, absorbance was measured for 10 minutes at 412nm to determine maximal enzymatic activity rate (V_max_). Where possible, values were then normalised to protein content (determined by Bradford method [Bio-Rad, UK]). For complex assays, the buffer reagents and spectrophotometer setting are given in Table 1. V_max_ was assessed over a ten-minute period at 37°C. Values were then either normalised to citrate synthase activity or protein content (determined by Bradford method [Bio-Rad, UK]). Enzymatic activity was calculated by using Equation 1.

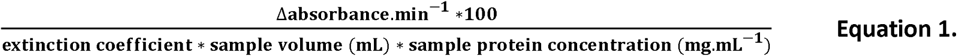

**Table 1.** Conditions for mitochondrial complex activity assays. λ - wavelength for the assay; ε - extinction coefficient; BSA - fatty acid–free bovine serum albumin; Cyt c - cytochrome c; Cyt c H_2_ - reduced cytochrome c; DCPIP - 2,6-dichlorophenolindophenol; DUB - decylubiquinone; DubH_2_ - decylubiquinol; KCN - potassium cyanide; KP - potassium phosphate buffer; Ub_1_ - ubiquinone_1_

|  | **CI** | **CII** | **CIII** | **CIV** |
| --- | --- | --- | --- | --- |
| **$\lambda$ (nm)** | 340 | 600 | 550 | 550 |
| **$\epsilon$ (mmol<sup>-1</sup> cm<sup>-1</sup>)** | 6.2 | 19.1 | 18.5 | 18.5 |
| **buffer** | KP (50 mM) | KP (25 mM) | KP (25 mM) | KP (50 mM) |
| **pH** | 7.5 | 7.5 | 7.5 | 7.0 |
| **Substrates/electron acceptors** | NADH, 100 μM<br>Ub<sub>1</sub>, 60 μM | Succinate, 20 mM<br>DCPIP, 80 μM<br>DUB, 50 μM | DubH<sub>2</sub>, 100 μM<br>Cyt c, 75 μM | Cyt c H<sub>2</sub>, 60 μM |
| **Detergent** | - | - | Tween-20<br>(0.025% (vol/vol)) | - |
| **Other reagent(s)** | BSA, 3 mg.mL<sup>-1</sup><br>KCN, 300 μM | BSA, 1 mg.mL<sup>-1</sup><br>KCN, 300 μM | KCN, 500 μM<br>EDTA, 100 μM | - |
| **Specific inhibitor** | Rotenone, 10 μM | Malonate, 10 mM | Antimycin A,<br>10 μg.mL<sup>-1</sup> | KCN, 300 μM |

### 2.8. In silico modelling

Oxygen-dependent mitochondrial respiration (v(O_2_)) was modelled using a two-population weighted sum Michaelis-Menten approach (Equation 2) to assess the effect of a mixed population of cytochrome c oxidase isoforms. Each COX4 isoform was assigned a distinct oxygen affinity (COX4I1, Κ_1_ = 0.058 kPa; COX4I2, K_2_ = 0.089 kPa).

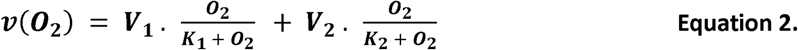

Where V_1_ and V_2_ are the fractional abundance of the COX4I1 and COX4I2 isoforms. Apparent P_50_ values for each fractional abundance ratio were calculated using the quadratic solution derived from the mixed Michaelis-Menten curve.

GSNO-dependent modulation of oxygen sensitivity was also modelled using experimentally derived GSNO dose-response data fitted with baseline-constrained Hill and linear models to the P_50_-GSNO dose response data. The fitted P_50_ values were subsequently used to scale oxygen affinity constants (K_1_ and K_2_ in equation 1) within the same two component Michaelis-Menten model described above (equation 1) to generate simulated respiration-oxygen relationships across a range of GSNO concentrations.

### 2.9. Statistics

All data analysis was performed with GraphPad Prism, version 9. All data points are shown, and data are presented as mean ± standard error of the mean (SEM). Data were tested for normal distribution using a Shapiro-Wilk test. Where data were not normally distributed, values were transformed by using log(Y) before statistical analysis^25^. When data were normally distributed, parametric paired (when CB and LV were isolated from the same animal) or unpaired t tests were used as indicated in the figure legends. In cases when data was not normally distributed and log transformation did not yield normal distribution; nonparametric tests were used (see figure legends). P-values are displayed in each graph, and statistical significance was set at p = < 0.05.

## 3. Results

### 3.1. Enhanced aerobic capacity but reduced oxidative phosphorylation efficiency in carotid body mitochondria

Citrate synthase activity (a common marker of tissue mitochondrial content) and abundance was lower in CB homogenates compared to the LV (Figure 1A & Supplementary Figure 1A), in line with findings for TOM20 protein abundance (Supplementary Figure 2A). However, when we measured aerobic capacity of mitochondria with high-resolution respirometry (example trace shown in Supplementary Figure 3A), CB respiration rates (normalised to citrate synthase) were higher than in the LV in LEAK_CI,_ OXPHOS_CI,_ OXPHOS_CI+CII_, ETC_CI+CII_ and ETC_CII_ states (Figure 1B-F). Complex IV respiration or 1-(L/P) were not different between the two tissues, however, RCR was higher in LV compared to CB samples (Figure 1G-I). Overall, these results suggest that CB tissue has a lower mitochondrial density and efficiency than the LV, but a greater aerobic capacity per mitochondria.

**Figure 1.**
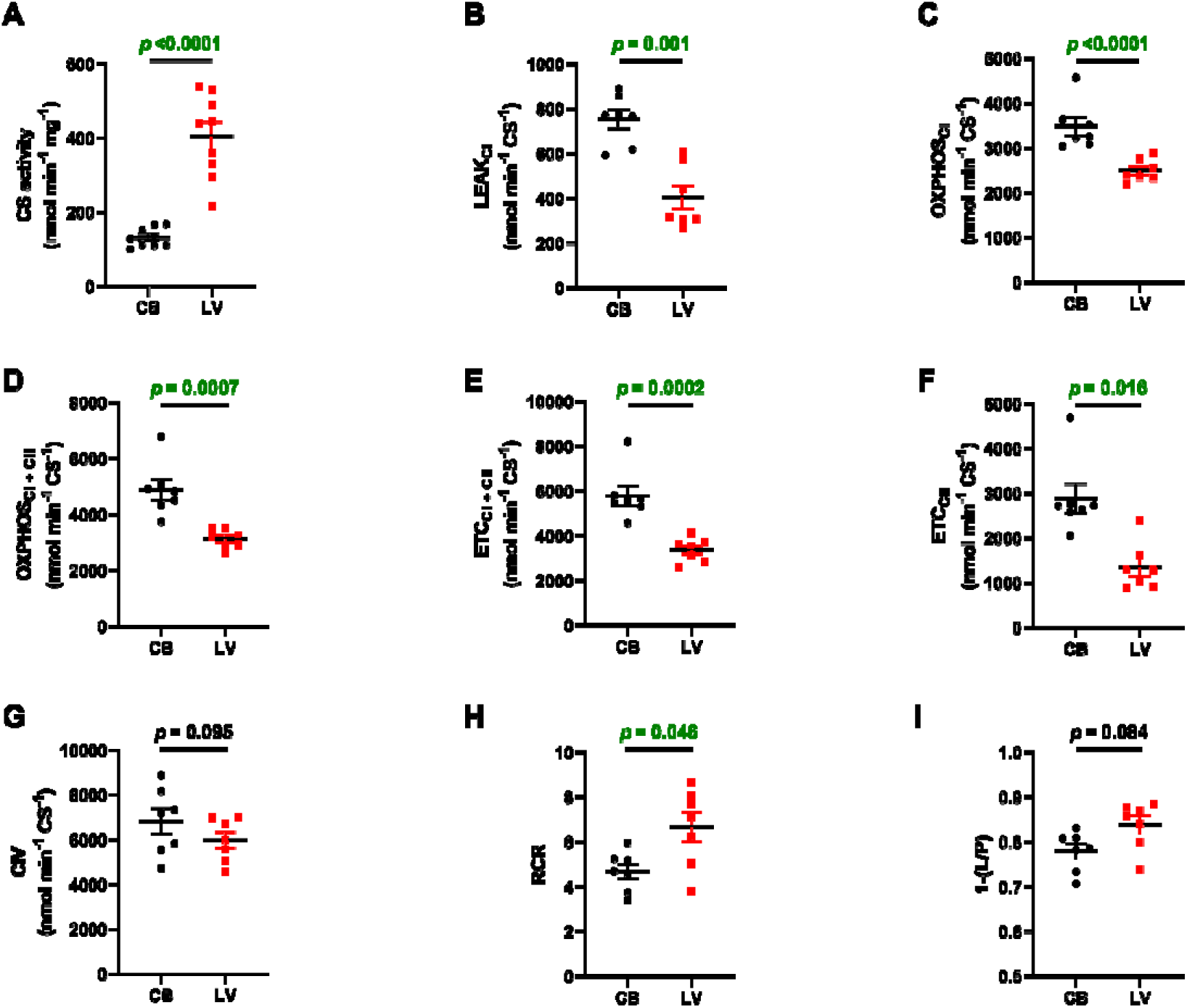
Comparison of oxygen consumption rates under different respiratory states between carotid body (CB) and left ventricle (LV) mitochondria. **A.** citrate synthase (CS) activity ( n = 9 in each group); **B.** LEAK respiration with complex I substrates (LEAK_CI_); **C.** mitochondrial respiration with complex I substrates (OXPHOS_CI_); **D.** OXPHOS with complex I and complex II substrates (OXPHOS_CI+CII_); **E.** maximum electron transfer capacity with complex I and complex II substrates (ETC_CI+CII_); **F.** ETC with complex II substates (ETC_CII_); **G.** maximum complex IV (CIV) oxygen consumption rate; **H.** respiratory control ratio (RCR); **I.** LEAK/OXPHOS coupling-control ratio. n = 7 in each group, two-tailed paired t-test (A-E & G-I) and two-tailed Wilcoxon test (F).

### 3.2. Carotid body have a high enzymatic activity, but low protein abundance of mitochondrial complexes

The higher aerobic capacity in CB vs LV isolated mitochondria could be explained by several factors, including a higher abundance or activity of mitochondrial complexes. Firstly, we performed mass spectrometry experiments to describe the subunit composition of each mitochondrial complex in the CB and LV mitochondria. Here, results show that all subunits detected are present in both tissues, but the abundance of most subunits is lower in the CB compared to the LV (Figure 2). No differences in abundance of NADH-ubiquinone oxidoreductase chain 4 (MT-ND4) and cytochrome c oxidase subunit 1 (MT-CO1) were found. However, the abundance of NADH dehydrogenase [ubiquinone] flavoprotein 3 (NDUFV3), cytochrome c oxidase subunit 7A2 (COX7A2), cytochrome c oxidase subunit 7A2-like (COX7A2L) and cytochrome c oxidase subunit 4 isoform 2 (COX4I2) were higher in the CB than LV. We validated these proteomics results with Western blots which also showed that the abundance of the individual mitochondrial complexes was lower in the CB compared to the LV, when normalised to either total protein (Supplementary Figure 1), or TOM20 (Supplementary Figure 2).

**Figure 2.**
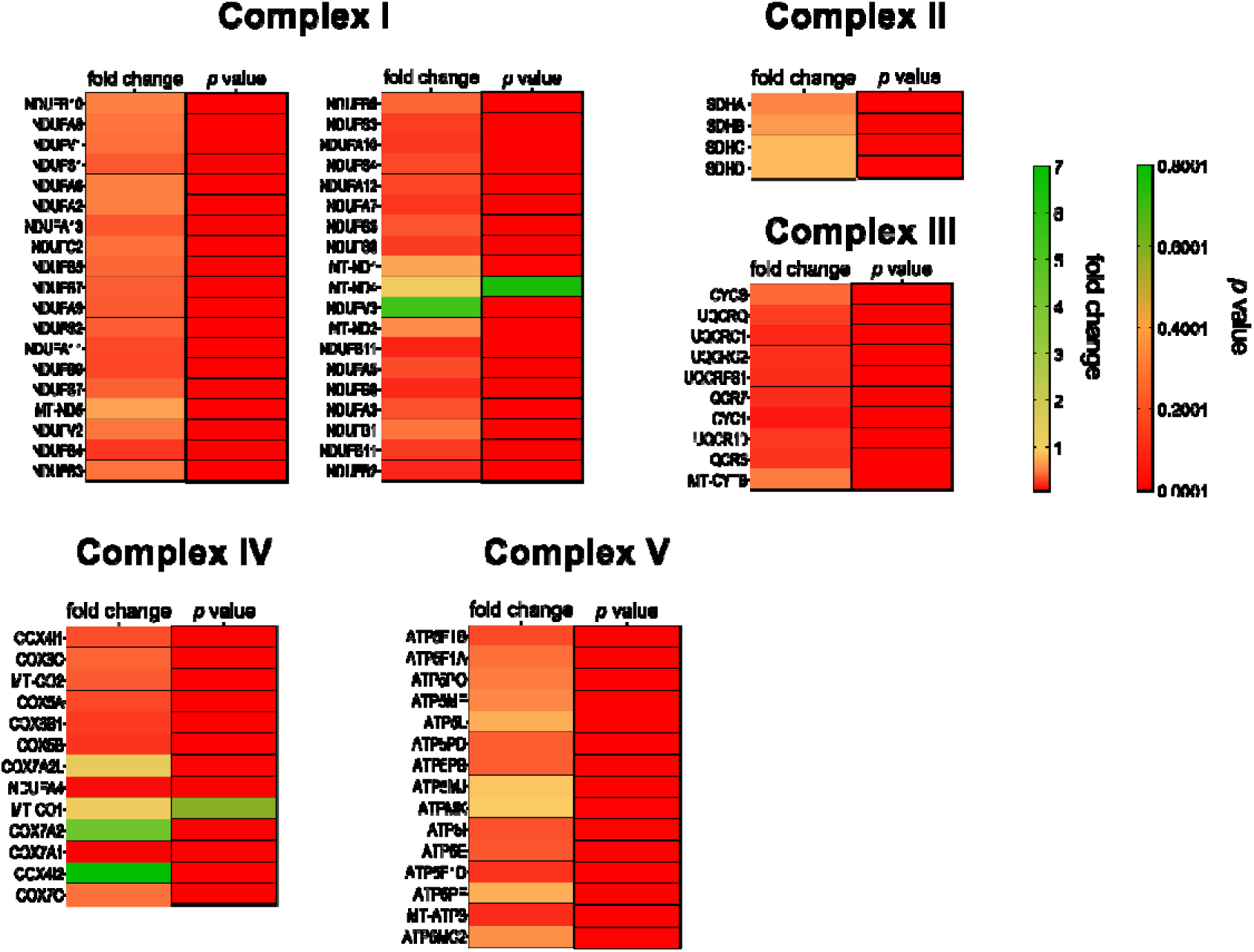
Differences in individual mitochondrial subunit abundance between carotid body and left ventricle. The majority of the detected subunits were lower in the carotid body compared to the left ventricle mitochondria. However, 4 subunits were higher in the carotid body: NADH dehydrogenase [ubiquinone] flavoprotein 3 (NDUFV3), cytochrome c oxidase subunit 7A2 (COX7A2), cytochrome c oxidase subunit 7A2-like (COX7A2L) and cytochrome c oxidase subunit 4 isoform 2 (COX4I2). Abundance of NADH-ubiquinone oxidoreductase chain 4 (MT-ND4) and cytochrome c oxidase subunit 1 (MT-CO1) was no different between the two samples. n = 12 in each group, two-tailed unpaired t-test.

Enzymatic activity of complexes I, II III and IV was also lower in the CB, when normalised to the total protein content (Figure 3A). However, when normalised to citrate synthase activity, as proxy for mitochondrial content, complex II, III and IV enzymatic activities were higher in the CB compared to the LV (Figure 3B.ii-iv). We conclude that mitochondrial content in LV is higher than CB, but importantly that enzymatic activity of complexes II, III and IV per mitochondria are higher in the CB.

**Figure 3.**
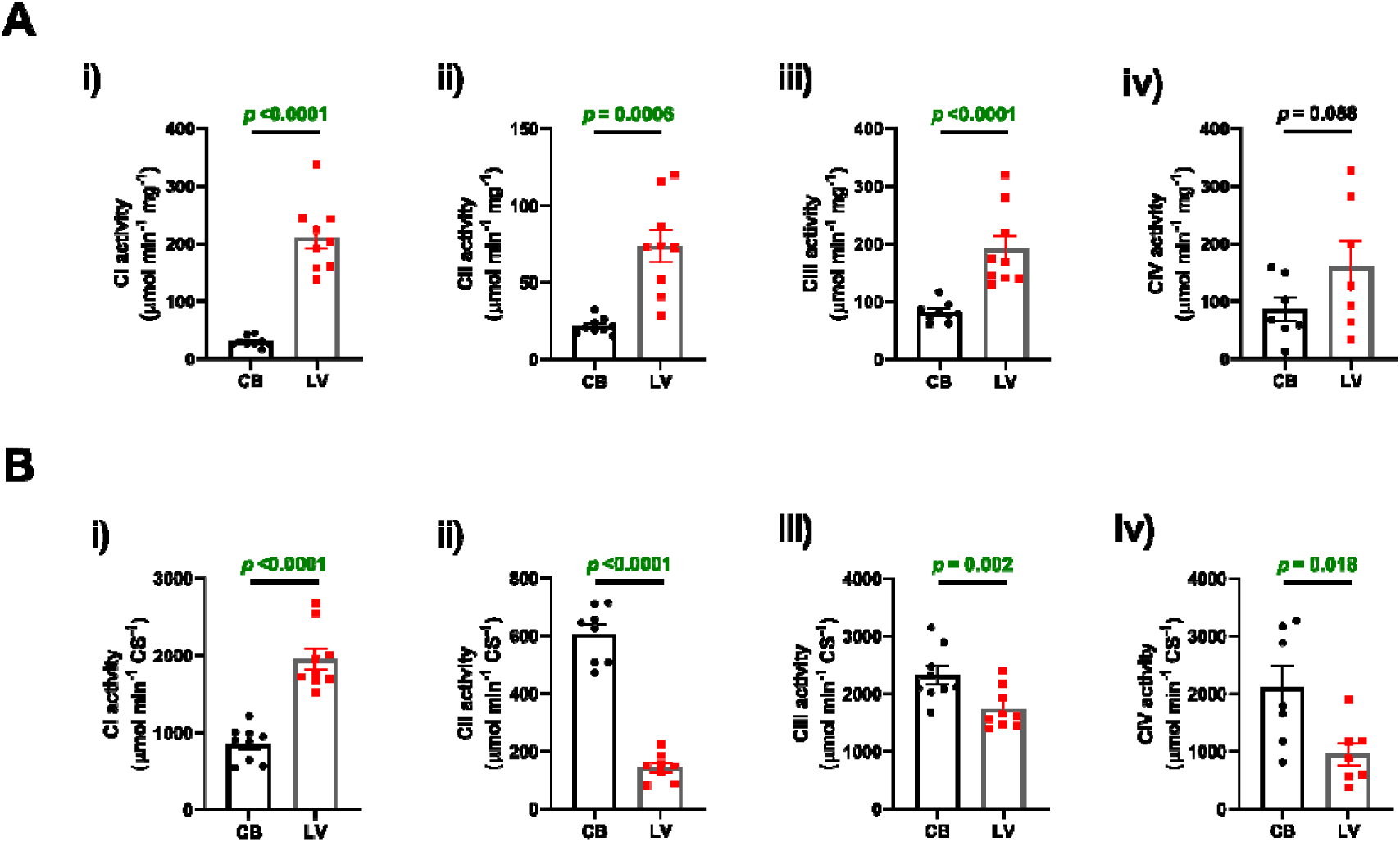
Higher mitochondrial complex activity in carotid body homogenates is revealed when activity is normalised to a mitochondrial marker. **A.** enzymatic activity normalised to the protein content and **B.** enzymatic activity normalised to citrate synthase activity; i) complex I (CI; n = 9 in each group); ii) complex II (CII; n = 9 in each group); iii) complex III (CIII; n = 9 in each group) and iv) complex IV (CIV; n = 7 in each group), two-tailed paired t-test.

### 3.3. Oxygen consumption rate determines H_2_O_2_ production in CB and LV mitochondria

In the CB, ROS production has been proposed to mediate signal transduction from the mitochondria to the cell membrane^4^, therefore, to characterise sites of ROS production, we measured H_2_O_2_ production under different respiratory states (example trace in Supplementary Figure 3B). When normalised to citrate synthase activity, H_2_O_2_ production was higher in the CB during LEAK_CI_, OXPHOS_CI_ and ETC_CII_, but not OXPHOS_CI + CII_, ETC_CI + CII_ or with CIII inhibited compared to the LV (Figure 4A). However, this effect disappeared when H_2_O_2_ production was normalised to oxygen consumption (Figure 4B), suggesting that differences in H_2_O_2_ production between CB and LV are determined by respiration rate, rather than intrinsic differences in electron leak.

**Figure 4.**
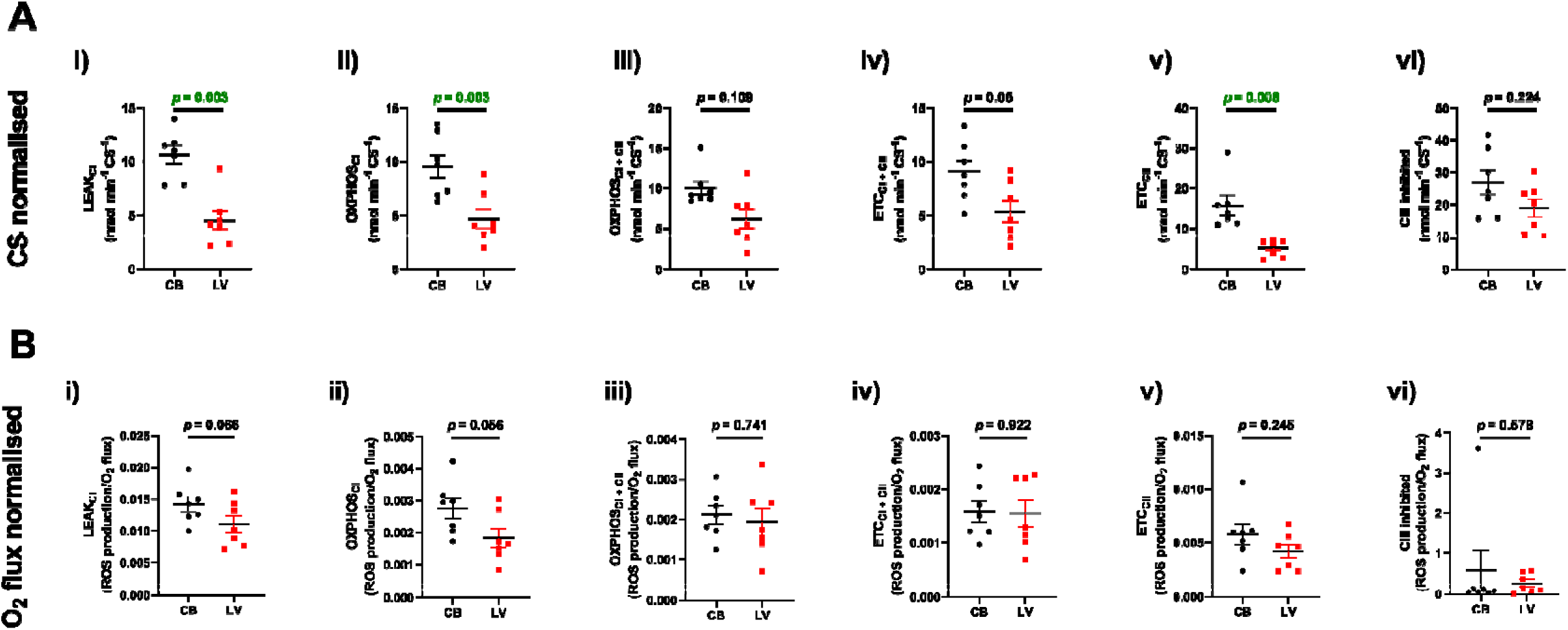
H_2_O_2_ production rate under different respiratory states is not different between the carotid body (CB) and left ventricle (LV) mitochondria when normalised to O_2_ flux. H_2_O_2_ production normalised to **A.** citrate synthase (CS) activity and **B.** oxygen (O_2_) flux. i) LEAK respiration with complex I substrates (LEAK_CI_); ii) mitochondrial respiration with complex I substrates (OXPHOS_CI_); iii) OXPHOS with complex I and complex II substrates (OXPHOS_CI+CII_); iv) maximum electron transfer capacity with complex I and complex II substrates (ETC_CI+CII_); v) ETC with complex II substrates (ETC_CII_); vi) with complex III being inhibited (CIII inhibited); n = 7 in each group, two-tailed paired t-test (A.i-A.ii & A.iv-A.v & B.i-B.iv) and two-tailed Wilcoxon test (A.iii & B.v).

### 3.4. High complex IV oxygen affinity, but a low RET capacity in carotid body mitochondria

Having characterised mitochondrial function in the CB, we next investigated COX subunit abundance and oxygen affinity in isolated mitochondria (see Supplementary Figure 4 for a representative experimental trace). First, we sought to confirm previous findings by Gao ^11^ that CB mitochondria had a higher abundance of the lower oxygen affinity complex IV subunit, COX4I2. Indeed, proteomic analysis revealed that the abundance of COX4I1 was higher in the LV mitochondria, but COX4I2 was higher in the CB (Figure 2 & Figure 5B-C). We also performed RNA in-situ hybridization analysis which showed that the expression of COX4I1 and COX4I2 RNA was co-localised within sheep CB type I cells (Figure 5A). When we measured the oxygen affinity of CB mitochondria with high-resolution respirometry, the complex IV P_50_ in CB mitochondria (0.089 ± 0.009 mmHg) was higher than in LV mitochondria (0.058 ± 0.01 mmHg). However, CB complex IV O_2_ sensitivity was well within the normal range for mammalian mitochondria. Furthermore, the complex IV P_50_ in CB mitochondria measured here is over 500 times lower than previous values reported in intact type I cells.

**Figure 5.**
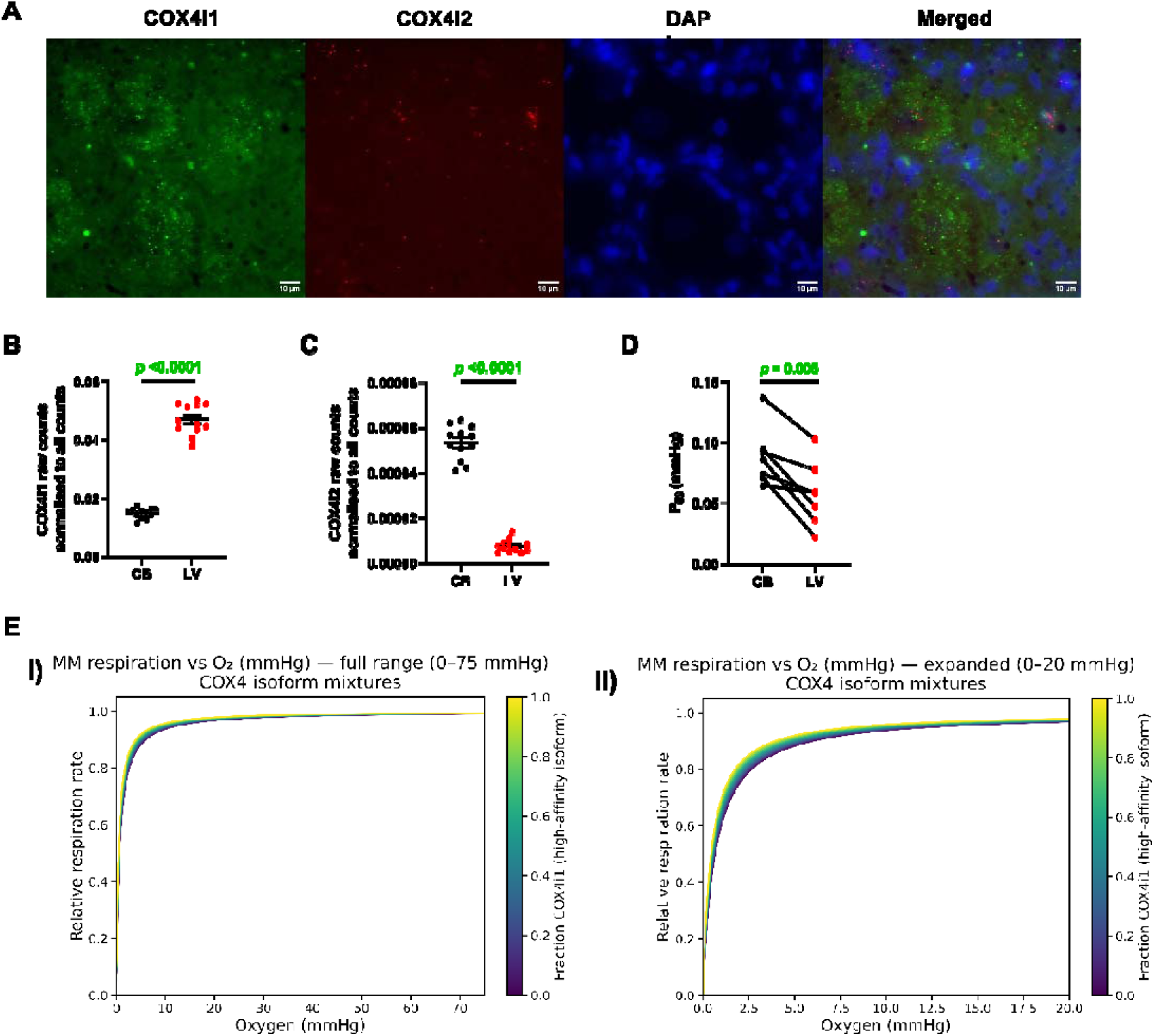
Carotid body (CB) cytochrome c oxidase has a high oxygen affinity despite higher abundance of COX4I2 subunit. **A.** expression of COX4I1 (green) and COX4I2 (red) RNA, counterstained with DAPI (nuclear stain, blue); abundance of **B.** COX4I1 and **C.** COX4I2 in the CB and LV mitochondria (n = 12 in each group, two-tailed unpaired t-test); **D.** 50% enzyme saturation value (P_50_) in isolated mitochondria (n = 7 in each group two-tailed paired t-test); **E.i**) relationship between relative oxygen consumption rate and oxygen tension at different fractions of COX4I1; **E.ii**) expanded version of the graph E.i.

It is important to note that mitochondria in this study were isolated from the whole CB, which contains both type I and type II cells. Oxygen sensing in the CB is restricted to type I cells, whereas type II cells exhibit a higher (i.e., more typical) oxygen affinity. Therefore, the inclusion of mitochondria from type II cells in our preparation could, in principle, shift the apparent P_50_ to lower values. However, given that mitochondria are present in higher numbers in type I cells (with only a few mitochondria present in type II cells; Supplementary Figure 5) and the fact that there are 3-5 type I cells per 1 type II cell, the contribution of type II mitochondria to the composite P_50_ is expected to be relatively minor. To address this, we performed computational modelling to estimate the expected P_50_ of a mixed mitochondrial population across a range of possible fractions of the high-affinity COX isoform. Within the constraints of the upper and lower oxygen affinities for the two COX4I isoforms, it is clear that altering the ratio of high to low affinity isoform within the tissue has very little impact on the P_50_ value and would be insufficient to confer physiological oxygen sensing capacity. Therefore, despite the higher oxygen affinity of type II cell mitochondria, their markedly lower cell content and mitochondrial density in the CB means that the overall mitochondrial pool and oxygen affinity is dominated by type I cells.

The monophasic nature of the oxygen consumption response further supports the notion that the oxygen affinity is high in the CB mitochondria (Supplementary Figure 6). If presence of COX4I2 subunit had a significant effect on overall oxygen affinity, a biphasic profile would be expected, with an initial decline at oxygen tensions corresponding to the lower affinity COX4I2 subunit (which is more abundant in type I cell mitochondria), followed by a second, steeper decline at lower oxygen tensions reflecting COX4I1 subunit (present in both type I and type II cell mitochondria). Instead, we observe a monophasic response, suggesting that COX4I2 has a relatively minor effect on overall measured oxygen affinity. Indeed, we do observe a slightly decreased oxygen affinity in the CB compared to the LV mitochondria.

Given that ROS derived from RET has been proposed to be involved in the CB response to hypoxia, we first evaluated RET capacity in the presence of succinate alone (example trace in Supplementary Figure 7). When normalised to citrate synthase activity, H_2_O_2_ production at baseline and during RET was higher in the CB compared to the LV (Figure 6A). At baseline, a similar relationship was observed when normalised to O_2_ flux, however, during RET, H_2_O_2_ production was lower in the CB mitochondria compared to the LV (Figure 6B) potentially suggesting that it may not have a crucial role in the oxygen sensing cascade.

**Figure 6.**
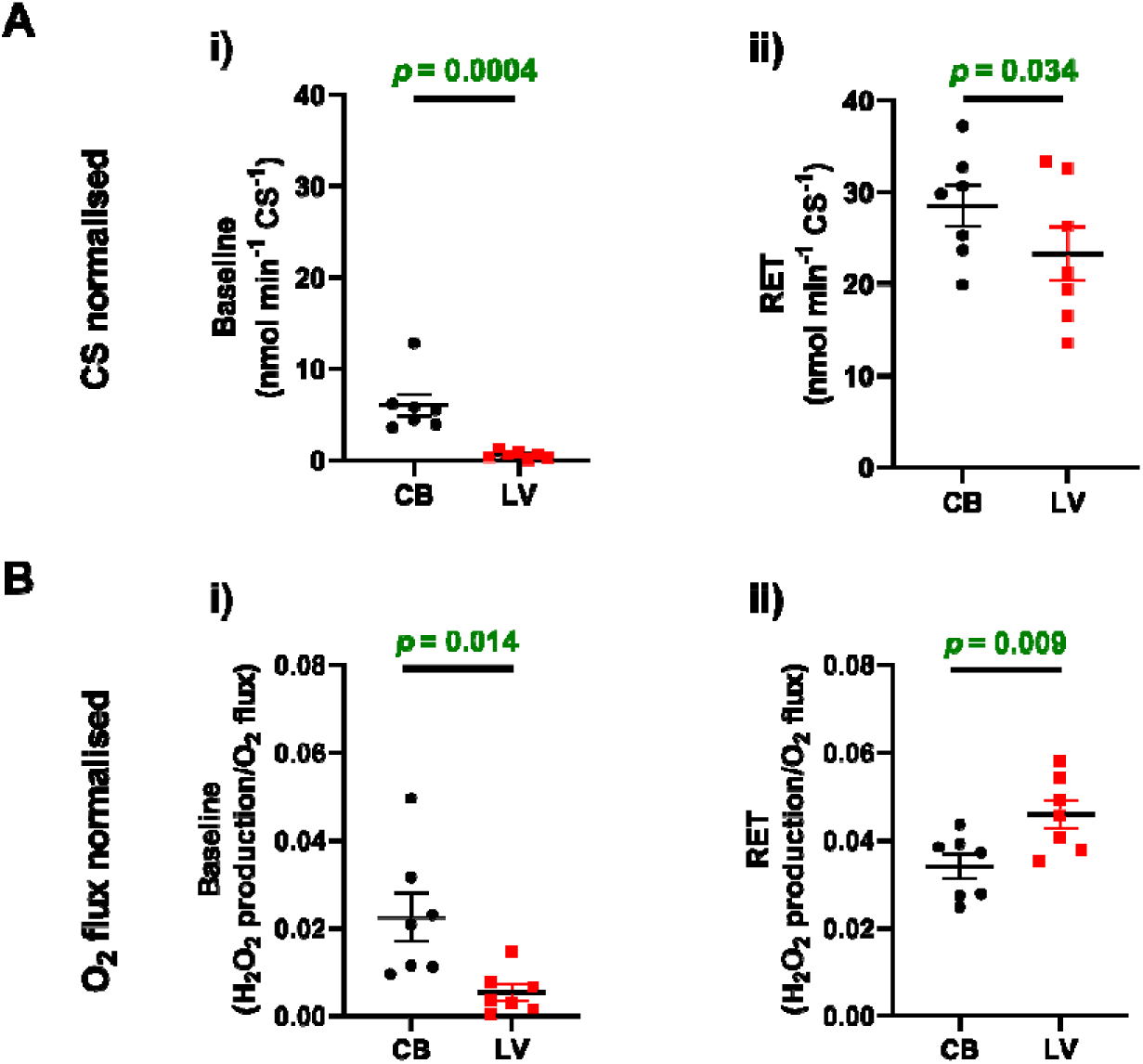
H_2_O_2_ production is higher at baseline and during reverse electron transport (RET) in carotid body (CB) mitochondria, however the increase in production rate is greater in the left ventricle (LV). H_2_O_2_ production normalised to **A.** citrate synthase (CS) activity and **B.** oxygen (O_2_) flux. i) baseline and ii) RET H_2_O_2_ production rate; n = 7 in each group, two-tailed paired t-test.

### 3.5. No evidence for unusual modulation of CB mitochondria through nitric oxide dependent pathways

The mitochondria-to-membrane hypothesis is based on the observation that type I cell oxygen affinity (and by proxy, apparent complex IV oxygen affinity) is unusually low compared to other tissue types. Having found no evidence to support this, we next examined whether known modulators of mitochondrial activity exert enhanced effects in CB mitochondria. NO is produced by nitric oxide synthase (NOS), which is widely expressed in the CB (Supplementary Figure 8). Using the NO donor GSNO (see Supplementary Figure 9 for representative trace), we assessed the ability of NO to directly inhibit complex IV via binding to its heme group. GSNO inhibited respiration in both tissue types in a concentration-dependent manner (Figure 7A & Supplementary Figure 10). This effect was unaffected by DTT but reversed by Hb(O₂)₄, consistent with a direct NO-mediated mechanism (Figure 7B–C).

**Figure 7.**
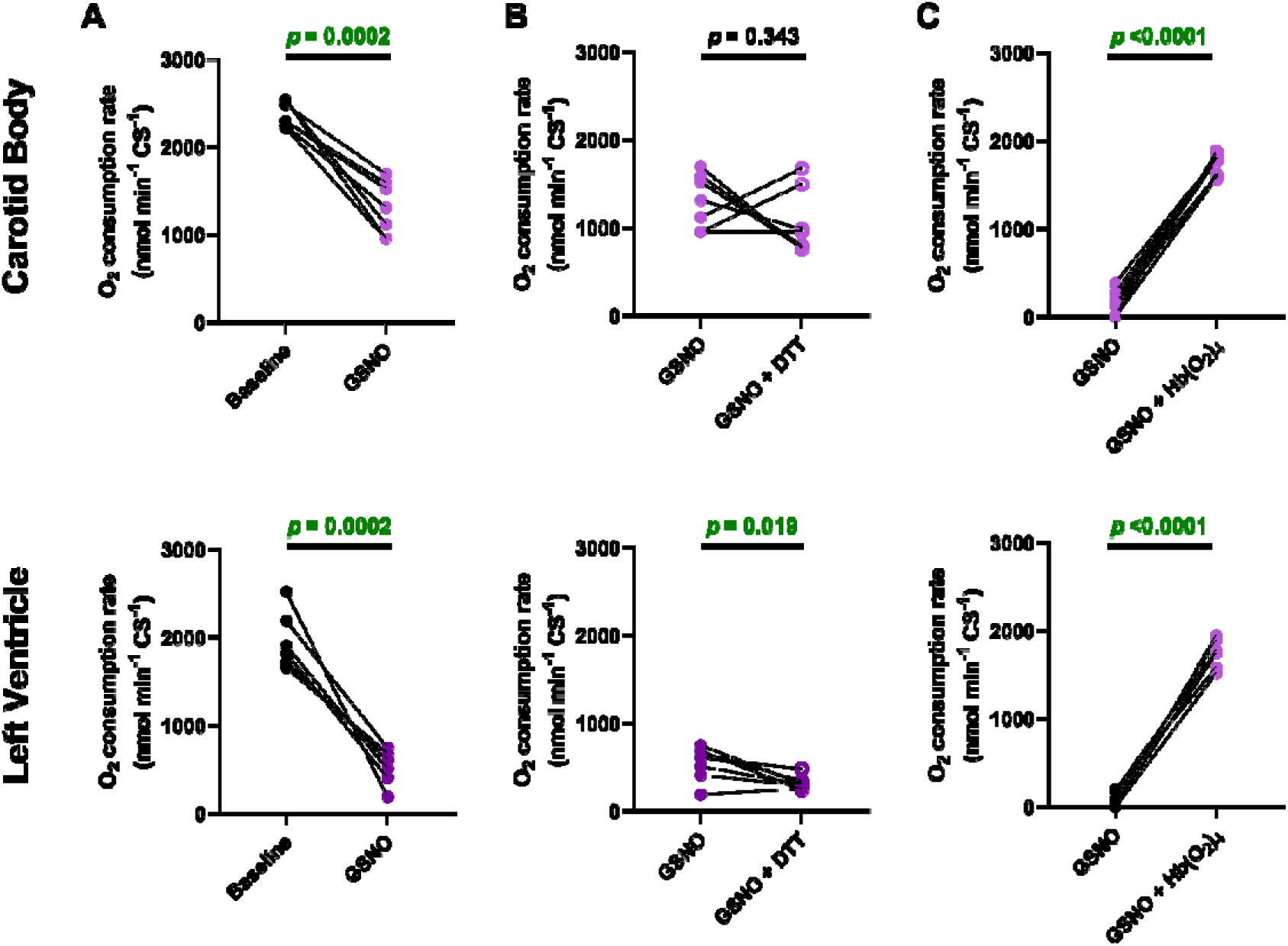
Nitric oxide (NO) inhibits mitochondrial respiration in carotid body (top panel) and left ventricle (bottom panel) mitochondria. The oxygen consumption rates in isolated mitochondria **A.** before and after addition of GNSO (n = 7 in each group); **B.** addition of DTT – to reverse S-nitrosylation (n = 7 in each group) and **C.** addition of oxyhaemoglobin (Hb(O_2_)_4_) to competitively reverse the effects of NO on complex IV (CB = 8, LV = 6). GSNO - S-Nitrosoglutathione; DTT - DL-Dithiothreitol; two-tailed paired t-test.

We then used the mitochondrially targeted S-nitrosylating agent mSNO to assess indirect NO-mediated effects via S-nitrosylation of mitochondrial complexes^26^. mSNO inhibited mitochondrial respiration in both tissues (Figure 8A & Supplementary Figure 11), whereas the mitochondrially targeted non-nitrosylating analog, mNAP, had only a minor effect (Supplementary Figure 12). The effects of mSNO were at least partially reversed by the addition of DTT, whereas, Hb(O_2_)_4_ had either minimal (LV) or no effect (CB), consistent with mSNO mediating its effects via S-nitrosylation (Figure 8B-C).

**Figure 8.**
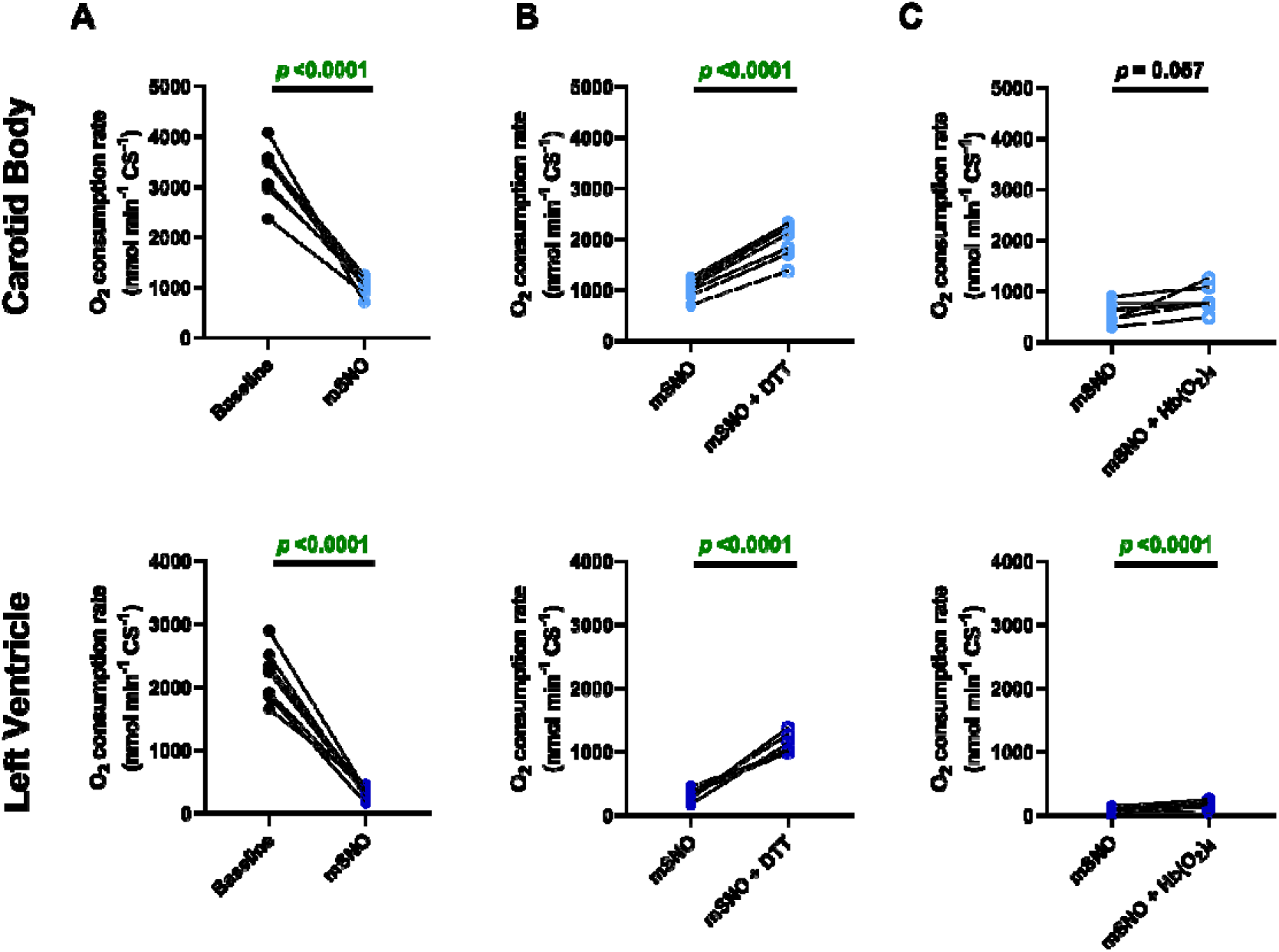
S-nitrosylation inhibits mitochondrial respiration in carotid body (top panel) and left ventricle (bottom panel) mitochondria. Oxygen consumption rates in isolated mitochondria **A.** before and after addition of mSNO, **B.** addition of DTT to reverse any effects of S-nitrosylation and C. reversal of the effects of NO with oxyheamoglobin (Hb(O_2_)_4_). mSNO – mitoSNO chloride; DTT - DL-Dithiothreitol; n = 7 in each group, two-tailed paired t-test.

Importantly, the effects of GSNO and mSNO on respiration were greater in LV mitochondria compared to the CB mitochondria (Supplementary Figure 13). Overall, these results suggest that NO can modulate CB complex IV activity via direct inhibition or through S-nitrosylation of mitochondrial complexes, although the magnitude of this effect is not unusual for mammalian mitochondria.

## 4. Discussion

In this study, we show that the oxygen affinity of complex IV in isolated CB mitochondria falls within the normal range for mammalian mitochondria. Thus, contrary to previous hypotheses, our data indicate that complex IV alone is unlikely to account for oxygen sensing in the CB. Instead, we suggest that additional signalling mechanisms modulate mitochondrial function in response to hypoxia. Consistent with this notion, NO suppressed respiration via both direct inhibition and S-nitrosylation, but these effects were not different from those observed in LV mitochondria. RET-associated ROS production was modestly elevated in CB mitochondria, potentially suggesting that other processes are involved in mediating signalling between mitochondria and cell membrane. Overall mitochondrial properties of CB were broadly similar to those of non-oxygen-sensing tissues.

### 4.1. Oxygen sensitivity of complex IV in the carotid body mitochondria

CB mitochondria have been widely described as unique in the literature, mainly due to their complex IV subunit composition and unusually low apparent oxygen affinity (∼46 mmHg) of intact CB type I cells ^4,9,11,27^. However, to date experiments have been performed on ex vivo CB preparations or isolated type I cells. It is well established that oxygen tension is much higher at the cellular interface (∼ 20 - 30 mmHg), than within the mitochondrial matrix (∼ 1 – 2 mmHg)^7,8,28-30^, arising from both oxygen diffusion limitations and mitochondrial respiration. Importantly, arterio-venous oxygen differences are much less in the CB than other tissues (less than 1 mmHg)^31^ due to a very high blood flow through the tissue. As such, the type I cell intracellular pO_2_ are likely greater than other tissues^31^. Therefore, previous approaches did not directly measure complex IV oxygen affinity, but rather assessed overall cellular oxygen affinity, with the unusually low values attributed to high expression of COX4I2^11^. Here we also show that the abundance of COX4I2 is indeed higher in the CB, and although complex IV P_50_ (0.089 ± 0.009 mmHg) was higher than LV mitochondria (0.058 ± 0.01 mmHg), it is well within the normal range (<0.5 mmHg) for mammalian mitochondria^7,8^, and over 500 times higher than expected from data on type I cell P_50_. While we acknowledge that the CB mitochondrial population was derived from a mixed population of type I and type II cells, computational modelling showed that the expected P_50_ of the mixed preparation would range from 0.063 to 0.081 mmHg with a fraction of COX4I1 isoform decreasing from 80 to 20 % of the total. Our measured CB mitochondrial P_50_ (0.089) is very close to this estimate, suggesting that intrinsic mitochondrial oxygen affinity is unlikely to be the principal determinant of CB oxygen sensitivity. The question then remains, if CB mitochondrial oxygen affinity is fairly normal, could other signalling mechanisms be modulating complex IV in intact CB cells?

### 4.2. NO as a modulator of oxygen affinity

NO decreases mitochondrial respiration through competitive binding of complex IV^15^, and S-nitrosylation of mitochondrial complexes. Consistent with this, we observed a concentration-dependent inhibition of respiration in CB mitochondria. However, this sensitivity was not markedly different from that of other tissues, including the LV.

Previous reports suggest the NO interaction with complex IV is different depending on the oxygen levels available^15^. When complex IV is predominantly oxidised (high oxygen availability) NO binds to the complex and is consumed (NO ^-^ formed, oxygen consumption unaffected)^15^. However, when complex IV is predominantly reduced (hypoxia), NO has an inhibitory effect which results in a decrease in NO metabolism and respiration rate but an increase in ROS signalling^15^. Complex IV also has a modulatory effect on NO production, namely when in its oxidised state, it limits NO production^32^. NO production is also dependent on oxygen concentration. Previous research showed that although the NOS activity and therefore the rate of NO production are dependent on oxygen level, the highest NO concentration in cultured cells was found under hypoxic conditions, specifically at 5 – 8% oxygen^33^. Importantly, an increase in NO production by NOS1 and NOS3 was also observed during hypoxia-ischaemia periods in the brain^34,35^. Therefore, it is plausible that under hypoxic conditions when complex IV changes its redox state, NO is produced in higher quantities by NOS present in the CB, and this leads to inhibition of complex IV and potentially an increase in ROS production and its signalling to the cell membrane. The process of inhibition of complex IV by NO under hypoxic conditions could be further strengthened if there were ‘compartments’ within the type I cell which allow for localised action of NO on mitochondrial function and therefore ROS signalling which could result in an initiation of the chemotransduction cascade. Furthermore, the presence of such compartments would mean that relative oxygen tension and NO levels within it would allow for the oxygen-sensing cascade to be initiated and a prompt signal transmission to the type I cell membrane. Indeed, a previous study performed by Holmes et al.^19^ reported that NO donation led to a moderate increase in hypoxic sensitivity in an ex vivo CB preparation. It is therefore plausible that oxygen affinity of complex IV is relatively high, but under hypoxic conditions, it is modulated by NO, leading to the initiation of the oxygen-sensing cascade. However, it is important note here that both this study and study by Holmes et al^19^ used NO concentrations that are far higher than those found in tissues/cells^33,36^. Hence, the question remains: are changes in oxygen affinity elicited by a physiological concentration of NO enough to timely respond to physiological hypoxia? Further research is necessary to answer this question.

Previous research has shown that the mitochondrial function could be affected by other factors, including hydrogen sulphide (H_2_S). H_2_S is a gasotransmitter produced by cystathionine γ-lyase (CSE) and in the CB, it has been proposed to play a crucial role in the oxygen sensing^37^. Briefly, under normoxic conditions, haem oxygenase-2 produces carbon monoxide under normoxic conditions, which prevents protein kinase G-dependent phosphorylation of CSE^38^. In hypoxia, this inhibition is removed and therefore promotes H_2_S production and initiates the hypoxic chemotransduction cascade^39^. Finally, H_2_S has been shown to increase NADH autofluorescence and therefore indicated that H_2_S could affect the mitochondrial ATP production^40^. Whilst H_2_S effects on mitochondrial function were not explored in this study, further work is warranted to fully dissect out the relationship between complex IV and NO and/or H_2_S.

### 4.3. ROS as a mediator of signalling between mitochondria and type I cell membrane

In CB tissue, reactive oxygen species (ROS) have been proposed to mediate signal transduction from mitochondria to the type I cell membrane as part of the oxygen-sensing cascade^20,21,41^. However, most studies to date have focused on hypoxia-induced changes in ROS production. Here, we provide the first measurements of basal ROS production in CB mitochondria and find no evidence of an unusual profile. When normalised to O₂ consumption, ROS production was comparable to LV tissue, and although H₂O₂ production during RET was modestly elevated in CB mitochondria, the magnitude of this effect was not striking. These results align to other studies that showed antioxidants do not fully abolish hypoxic responsiveness^42-44^ with one study stating that ROS production in type I cells decreases in response to hypoxia^45^. Clearly, further studies are required to fully assess the role of ROS in CB mitochondria under hypoxic conditions. However, based on our findings, ROS production via RET is unlikely to serve as the primary signal linking mitochondria to the cell membrane in the chemotransduction cascade. Instead, other candidates, such as MgATP^14,46^ and AM(D)P:ATP ratio^47^ should be considered as potential candidates or that multiple signalling pathways are involved depending on the severity of hypoxia (for review see^5^).

### 4.4. Mitochondrial aerobic capacity in the carotid body

Despite the proposed crucial role of mitochondria in CB oxygen sensing, very little is known about its oxidative capacity. Here, we show that CB mitochondria have a higher intrinsic aerobic capacity than LV mitochondria, but a lower mitochondrial content and reduced efficiency of ATP production (lower RCR). The higher rates of respiration in CB mitochondria were supported by an increased enzymatic activity of mitochondrial complexes II, III and IV. In agreement with our findings, Daly ^31^ showed that the CB has higher blood flow through the tissue, and also a higher respiration rate compared to other tissues (as extrapolated from blood flow, arteriovenous oxygen difference and CB weight). Biscoe and Duchen also concluded that isolated intact type I cells consume high amounts of oxygen compared to dorsal root ganglion cells^10^, with the total rate of consumption being close to their maximal capacity^10^.

The higher citrate synthase activity and mitochondrial efficiency observed in LV compared to CB are not unexpected, given that mitochondria occupy up to one-third of total cell volume in cardiac myocytes to meet the high energetic demands of excitation–contraction coupling^48^. Previous work has shown CB mitochondria are small and oval in shape (as opposed to being elliptical or rod-shaped as in the heart), potentially reflecting that the CB is of a neuronal origin^49^. Indeed, neuronal tissues, such as the superior cervical ganglion, have small round mitochondria with a lower oxidative capacity than the heart^50^. The lower efficiency of CB mitochondria was associated with a higher LEAK respiration rate, which suggests CB mitochondria may have a high proton leak. While an increased proton leak reduces ATP capacity, it is proposed to have protective effects by reducing ROS production (‘mild uncoupling’)^51,52^.

A previous study using mouse CB showed a high concentration of succinate in homogenate samples in comparison to brain and adrenal medulla samples, suggesting that succinate may be an important substrate for mitochondrial respiration^20^. Our data supports these finding as succinate dehydrogenase activity is higher in CB mitochondria. High succinate metabolism may be then further supported (or matched) by complex III and complex IV which activities are also higher in the CB mitochondria.

### 4.5. Limitations

The primary limitation of this study is that mitochondria were isolated from the whole CB, resulting in a heterogeneous population of mitochondria from type I and type II cells. This means we are unable to directly attribute the measured properties to mitochondria from type I cells alone. However, as supported by our computational modelling, the impact of this heterogeneity on the results is expected to be minimal, as the overall mitochondrial pool is predominantly derived from type I cells. Another limitation is that experiments were performed under fully oxygenated conditions; therefore, the effects of hypoxia on these parameters were not assessed and represent an important avenue for future research. Finally, this study focused primarily on mitochondrial composition and function in the CB using isolated mitochondria, which reveals their inherent properties but does not capture how these processes operate within intact type I cells. As such, we have not explored how NO and/or S-nitrosylation influence type I cell function, neurotransmitter release, or carotid sinus nerve activity, particularly under hypoxic stimulation.

## 5. Conclusions

Despite the fact that CB mitochondria have a unique composition of complex IV which could result in an unusually lower oxygen affinity, our data show that the oxygen affinity of complex IV in CB tissue lies within the normal range for mammalian mitochondria. Therefore, we conclude that complex IV is not the sole site of oxygen sensing in the carotid body and is more likely that complex IV function is modulated by additional factors to confer oxygen sensing properties to the CB. We found that NO inhibited mitochondrial respiration, suggesting it may contribute to the oxygen-sensing cascade. However, these effects were not enhanced or unique in CB mitochondria, and their precise role in oxygen sensing remains unclear. Additionally, RET-induced ROS production was elevated in the CB as expected, however overall lower than in the LV, signifying that other processes may be involved in the oxygen chemotransduction cascade. Clearly, further studies are needed to fully elucidate the mechanisms underlying oxygen sensing in the carotid body.

## CRediT authorship contribution statement

Agnieszka Swiderska: Conceptualisation, Data curation, Formal analysis, Investigation, Methodology, Validation, Visualization, Writing – original draft, Writing – review & editing. Michael P. Murphy: Conceptualisation, Supervision, Writing – review & editing. Gina L. J. Galli: Conceptualisation, Methodology, Supervision, Writing – original draft, Writing – review & editing. Andrew W. Trafford: Conceptualisation, Funding acquisition, Methodology, Supervision, Project administration, Writing –original draft, Writing – review & editing.

## Acknowledgements

We thank The University of Manchester Biological Mass Spectrometry core facility (RRID code: SCR_020987) for support with tissue proteomics sample preparation, data acquisition and analysis. The authors thank the staff in the EM Core (SCR_021147) Facility in the Faculty of Biology, Medicine and Health for their assistance, and the Wellcome Trust for equipment grant support to the EM Core Facility. Authors thank Dr Christian Pinali for his comments on the manuscript.

## Data availability statement

Data associated with this project will be made available on acceptance for publication on a CC-BY Creative Commons Attribution licence through The University of Manchester FigShare repository (DOI provided on acceptance).

