## Supplemental Material for "Carotid body mitochondria exhibit normal oxygen affinity despite COX4I2 enrichment"

**Supplement**

**Supplementary Methods**

**1. Western blot**

Western blot (as previously described^1^) was used to determine protein abundance of different mitochondrial proteins. Firstly, CB tissue was homogenised in RIPA buffer containing protease and phosphatase inhibitor cocktail (1:100 dilution) and protein concentration was determined by Bradford assay as per manufacturer’s recommendations (DC protein assay kit, Bio-Rad, UK). NuPAGE™ 4-12% Bis-Tris pre-cast polyacrylamide protein gel was used to run all samples following which separated proteins were transferred onto a nitrocellulose membrane (see Supplementary Table 1 for all details). Membranes were then incubated with a total protein stain and imaged before blocking with 5% milk. After blocking, primary antibodies were added to membrane and incubated overnight. Finally, membranes were incubated with the anti-rabbit secondary antibody and imaged. All blots were run in triplicate and averaged to obtain the final result. The abundance of the target protein was normalised to the total protein stain and internal control.

**2. Electron microscopy**

Carotid body tissue cubes (approx. 1 mm^3^) were prepared and immersion-fixed in Karnovsky's fixative. Samples were washed in water prior to staining. Staining was performed in two stages, firstly using 1% osmium tetroxide and 1.5% potassium ferrocyanide, followed by staining using 1% uranyl acetate (samples were washed in water after each step). Dehydration was performed using increasing concentration of ethanol washes (25, 50, 70, 90 and 3 washed in 100%), followed by further dehydration in pure acetone (3 washes). TAAB medium resin was used to infiltrate samples (incubation in 25, 50, 75 and 3x 100% resin mixed with acetone). Finally, samples were embedded and cured at 60°C for approximately 36 hours in pure resin.

80nm sections were cut with Leica UC7 ultramicrotome and examined with Thermo Fisher Talos L120C transmission electron microscope at 120kV acceleration voltage.

**3. Gene expression**

**3.1. RNA extraction**

RNA was extracted from whole CB. 50 mg of tissue was placed in 1 mL of TRIzol® (Invitrogen), incubated for 5 minutes on ice and then homogenised. 200 µL of chloroform was added, mixed with the rest of the solution and incubated for 2-3 minutes. Afterwards, each sample was centrifuged at 10,000 g for 15 minutes (4°C). The aqueous portion was transferred to a fresh Eppendorf and 100% ethanol was added (1:1.5 ratio). Following this, RNA was extracted using Qiagen miRNeasy Micro Kit (Qiagen) according to manufacturer’s instructions.

**3.2. RNA to cDNA conversion**

The amount of RNA was determined using a Nanodrop (ND-1000). 500ng RNA converted to cDNA using High-Capacity RNA-to-cDNA kit (Applied Biosystems) by following manufacturer’s instructions.

**3.3. Real Time qPCR**

Real time qPCR (RT-qPCR) was performed with Taqman Universal master mix (Applied Biosystems), on a QuantStudio Real-Time PCR machine (Applied Biosystems) by using differential heat and time cycles (1 cycle of 5 minutes at 50°C and 10 minutes at 95°C followed by 40 cycles of 15 seconds at 95°C and 1 minute at 60°C). Primers against NOS1 (Oa04714380_m1), NOS3 (ARKA4EA), PGK1 (Oa04657427_gH), RPLP0 (Oa04824512_g1). The comparative C_T_ method (ΔC_T_) was used to quantify the fold-expression of target genes. All C_T_ values were standardised against the two relative housekeeping genes C_t_ values (ΔΔC_T_) and then analysed using the formula 2- ΔΔC_T_.

**Supplementary Table 1.Summary of Western blot running conditions and reagent sources.**

CS – citrate synthase; NOS3 – endothelial nitric oxide synthase; TOM20 – translocase of outer mitochondrial membrane 20.

| **Protein of interest** | **Amount of protein loaded (µg)** | **Running conditions** | **Primary antibody** | **1° Ab.**  **Dilution** | **Secondary antibody** | **2° Ab.**  **Dilution** |
| --- | --- | --- | --- | --- | --- | --- |
| **NDUFS1 (complex I)** | **10** | **MES, 200 V, 50 minutes** | **ProteinTech, 12444-1-AP, Rabbit polyclonal** | **1:3000** | **IRDye® 800CW Donkey anti-Rabbit IgG Secondary Antibody (LI-COR Biosciences, UK)** | **1:20 000** |
| **SDHA (complex II)** | **10** | **MES, 200 V, 50 minutes** | **ProteinTech, 14865-1-AP, Rabbit polyclonal** | **1:2000** | **IRDye® 800CW Donkey anti-Rabbit IgG Secondary Antibody (LI-COR Biosciences, UK)** | **1:20 000** |
| **UQCRC2 (complex III)** | **10** | **MES, 200 V, 50 minutes** | **ProteinTech, 14742-1-AP, Rabbit polyclonal** | **1:4000** | **IRDye® 800CW Donkey anti-Rabbit IgG Secondary Antibody (LI-COR Biosciences, UK)** | **1:20 000** |
| **MTCO1 (complex IV)** | **10** | **MES, 200 V, 50 minutes** | **Abcam, ab14705, Mouse monoclonal** | **1:2000** | **IRDye® 800CW Donkey anti-Mouse IgG Secondary Antibody (LI-COR Biosciences, UK)** | **1:20 000** |
| **ATP5a (Complex V)** | **10** | **MES, 200 V, 50 minutes** | **Abcam, ab14748, Mouse monoclonal** | **1:1000** | **IRDye® 800CW Donkey anti-Mouse IgG Secondary Antibody (LI-COR Biosciences, UK)** | **1:20 000** |
| **TOM20** | **10** | **MES, 200 V, 50 minutes** | **ProteinTech, 11802-1-AP, Rabbit polyclonal** | **1:1000** | **IRDye® 800CW Donkey anti-Rabbit IgG Secondary Antibody (LI-COR Biosciences, UK)** | **1:20 000** |
| **CS** | **10** | **MOPS, 150V, 90 minutes** | **ProteinTech, 16131-1-AP, Rabbit polyclonal** | **1:2000** | **IRDye® 800CW Donkey anti-Rabbit IgG Secondary Antibody (LI-COR Biosciences, UK)** | **1:20 000** |
| **NOS3** | **10** | **MOPS, 150V, 90 minutes** | **BD Transduction Laboratories™, 610297, Mouse monoclonal** | **1:2500** | **IRDye® 800CW Donkey anti-Mouse IgG Secondary Antibody (LI-COR Biosciences, UK)** | **1:20 000** |

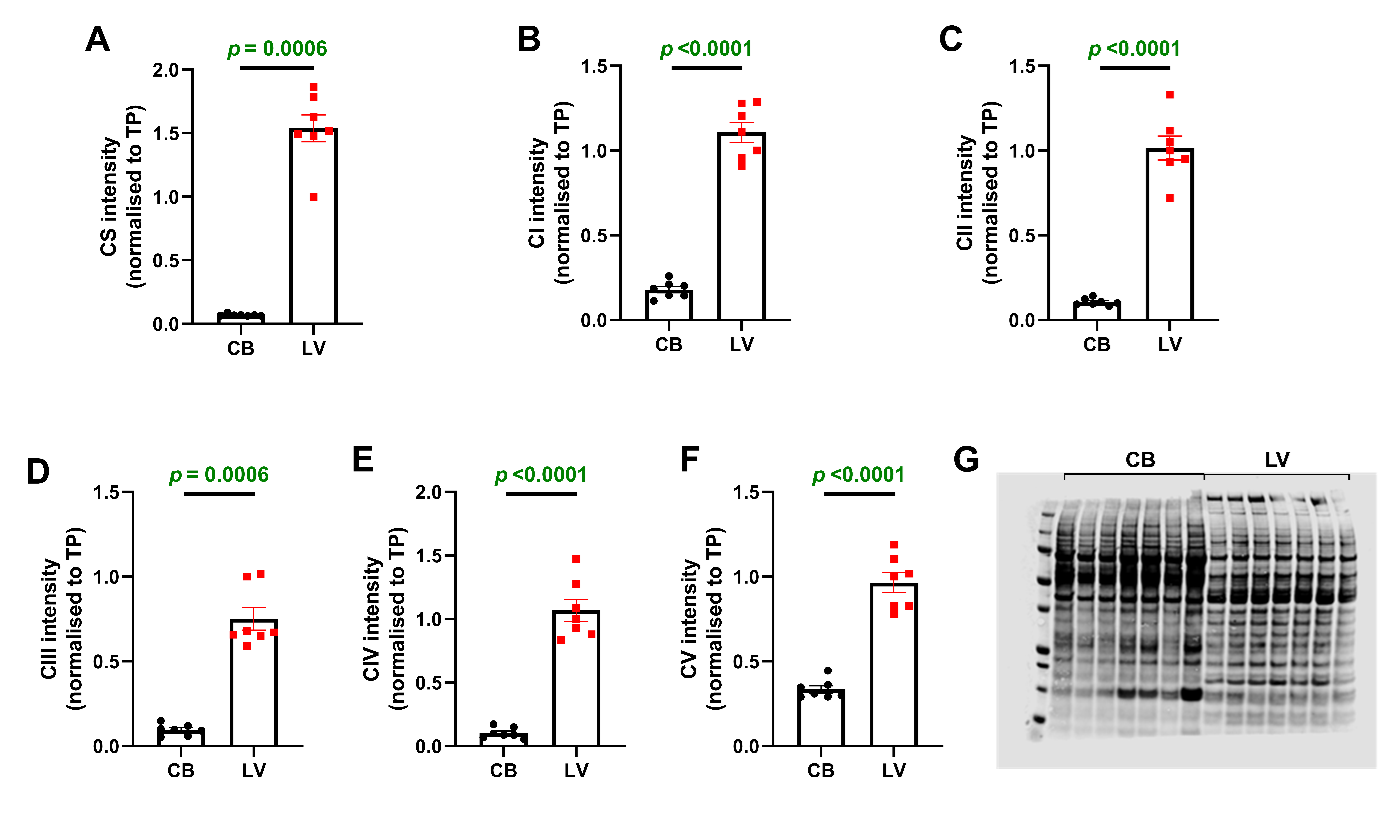

Supplementary Figure 1. Protein abundance of each individual mitochondrial complex is higher in the left ventricle (LV) compared to the carotid body (CB).

A. citrate synthase (CS); B. complex I (CI), **C**. complex II (CII), **D**. complex III (CIII), **E**. complex IV (CIV) and **F**. complex V (CV) protein abundance per total protein (TP), **G**. representative image of total protein stain; n = 7 in each group, average of 3 technical repeats. unpaired t-test (B-C & E-F), Mann-Whitney test (A & D).

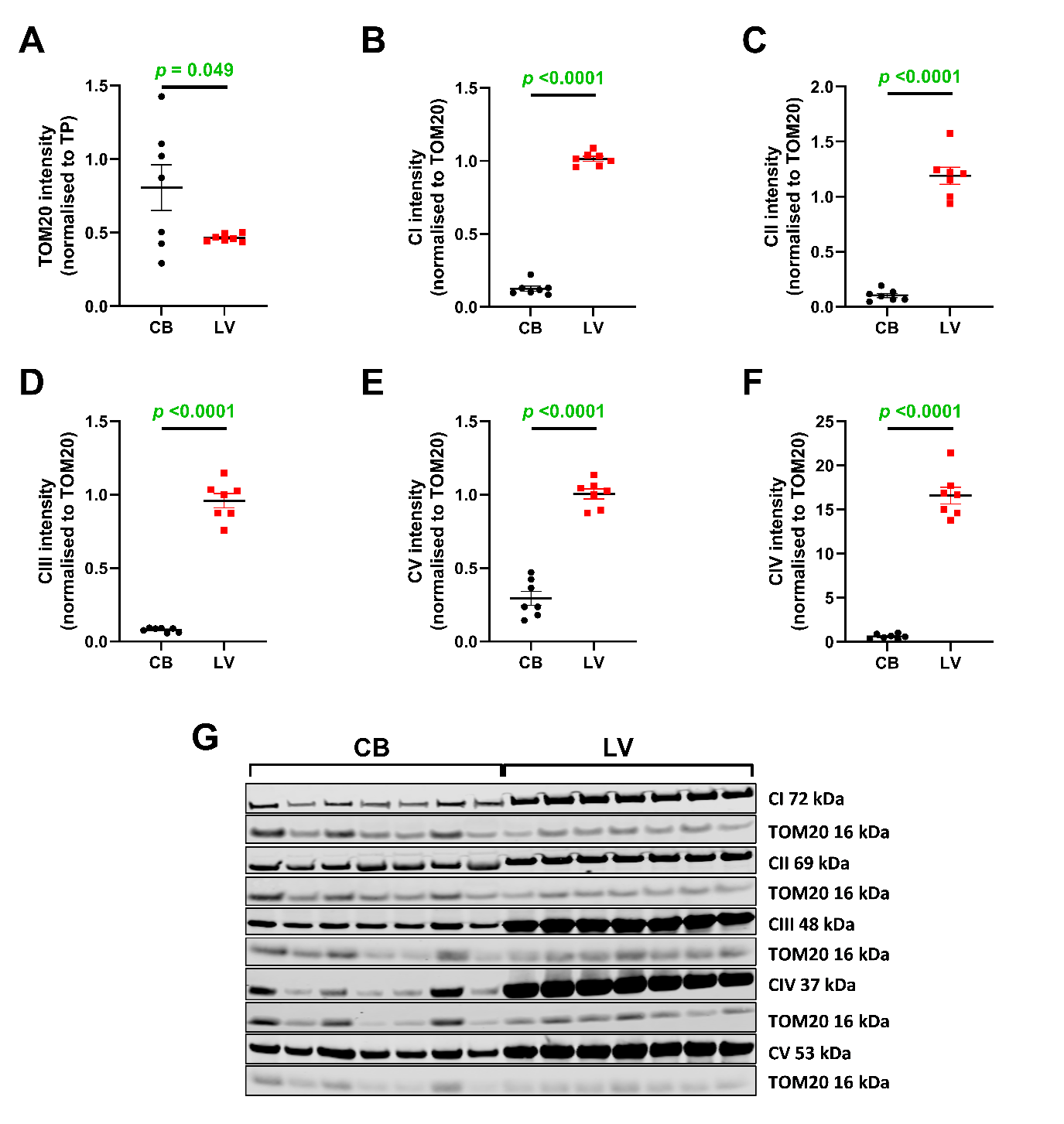

Supplementary Figure 2. **The protein abundance of each individual mitochondrial complex per housekeeper is statistically higher in the left ventricle (LV) compared to the carotid body (CB).**

Protein abundance of **A**. TOM20 (normalised to total protein (TP) and mitochondrial **B**. complex I (CI), **C**. complex II (CII), **D**. complex III (CIII), **E**. complex IV (CIV) and **F**. complex V (CV); **G**. representative western blot images; n = 7 in each group, average of 3 technical repeats, parametric unpaired t-test.

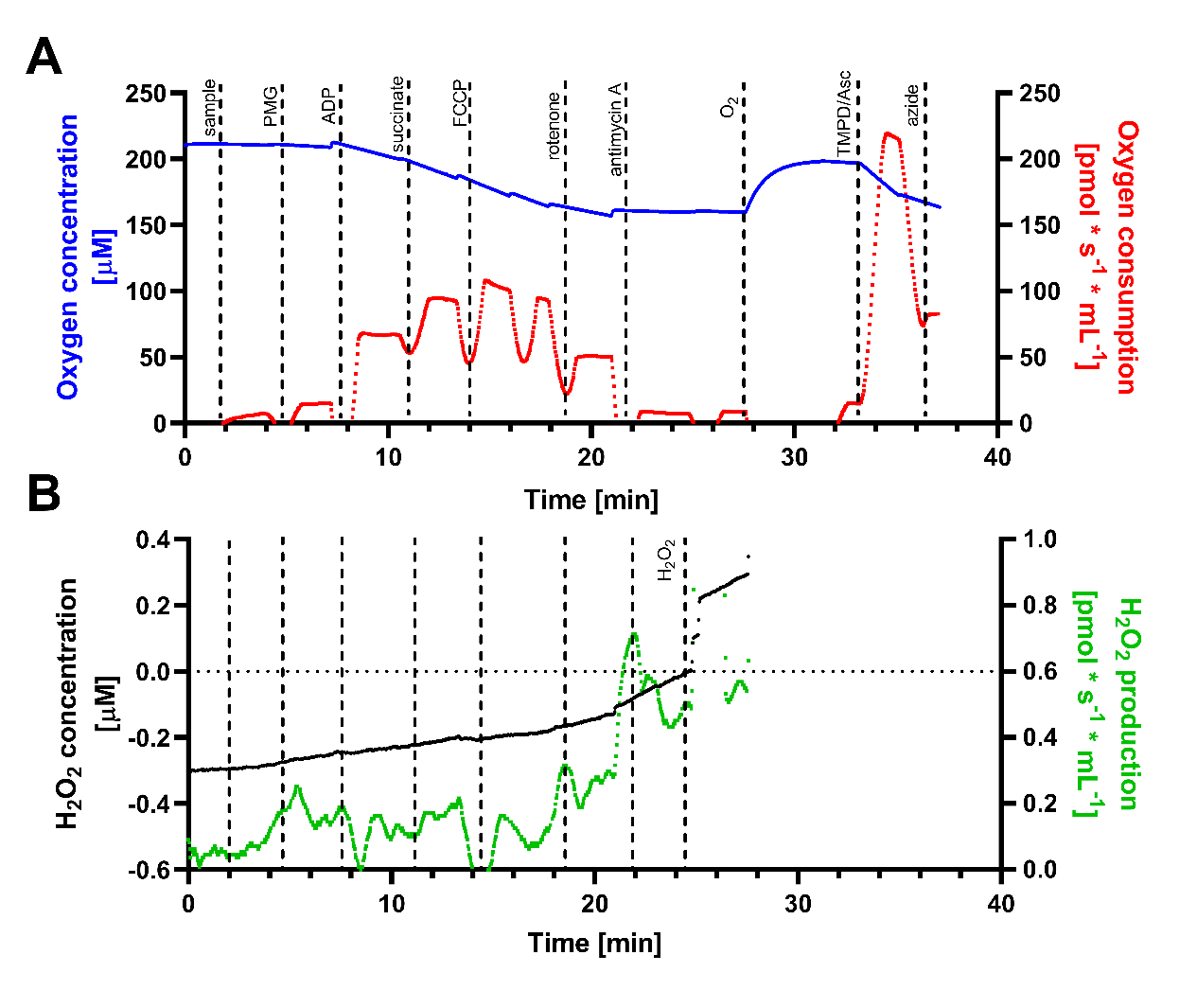

Supplementary Figure 3. Representative trace of substrate-uncoupler-inhibitor titration (SUIT) protocol.

Original traces of the SUIT protocol: A. mitochondrial oxygen consumption and B. H_2_O_2_ production. Prior to experimentation, Amplex UltraRed, horseradish peroxidase and superoxidase dismutase were added to measure the H_2_O_2_ production. Chemicals were added in order as indicated on traces to measure oxygen consumption rate and H_2_O_2_ production rate under different respiratory states. Note that measurements were made when the trace was stable and injection artefacts excluded. ADP - adenosine diphosphate, Asc – ascorbate, FCCP = carbonyl cyanide-4-(trifluoromethoxy)phenylhydrazone, PMG – pyruvate, malate, glutamate, TMPD = N,N,N,N-tetramethyl-p-phenylenediamine.

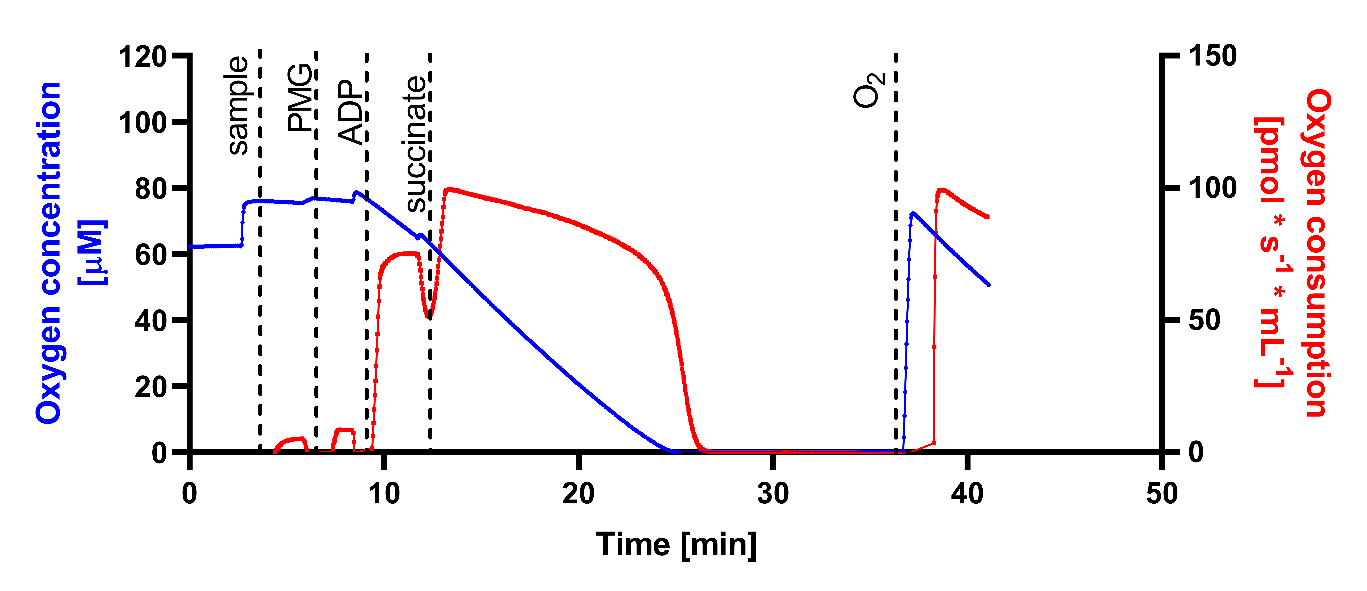

Supplementary Figure 4. Representative trace of the oxygen affinity protocol.

ADP - adenosine 5′-diphosphate, PMG – pyruvate, malate, glutamate.

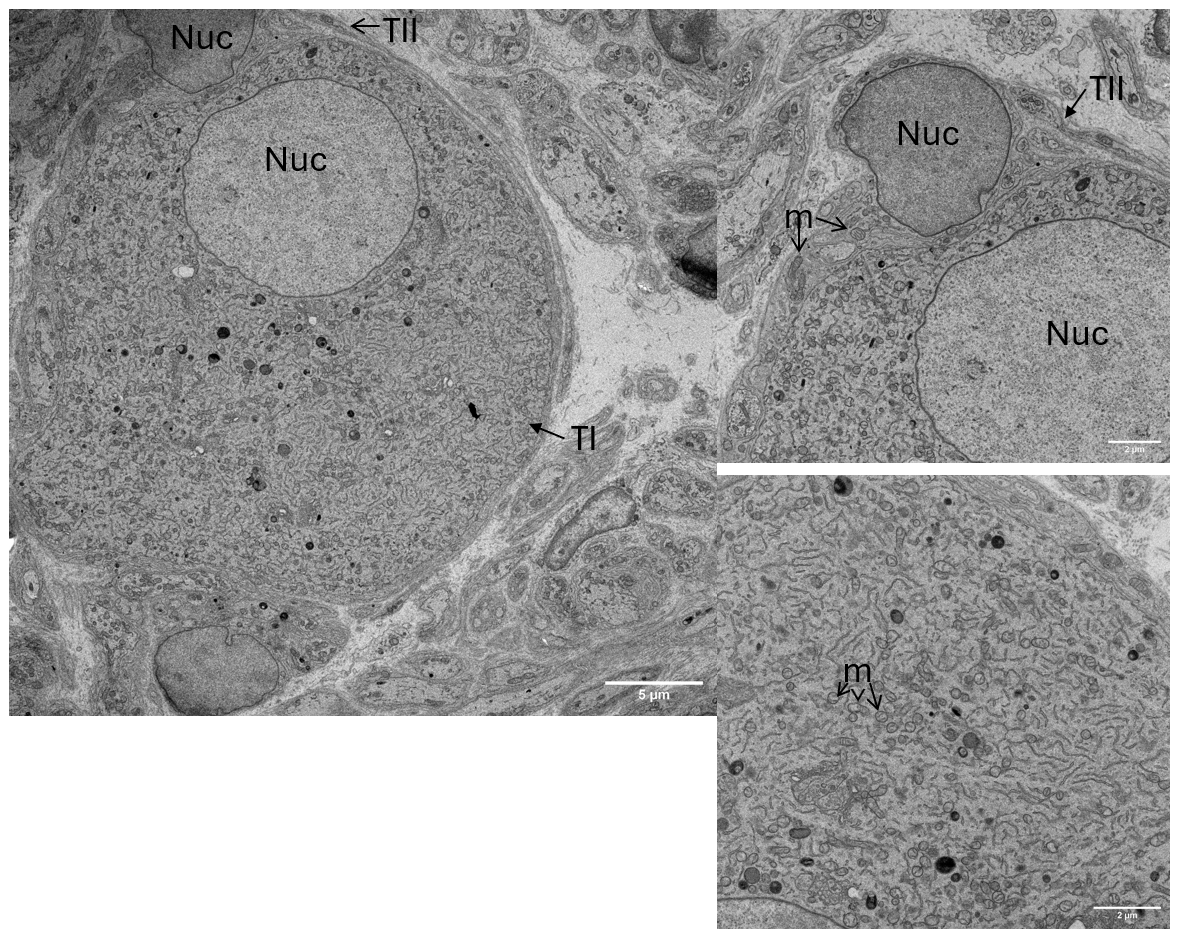

**Supplementary Figure 5. A representative electron microscopy image of carotid body cells.**
Image shows an oval type I (TI) cell with an associated type II (TII) cell. It is visible that while TI cell contains many mitochondria (m), there is far less mitochondria present in TII cell. Nuc – nucleus.

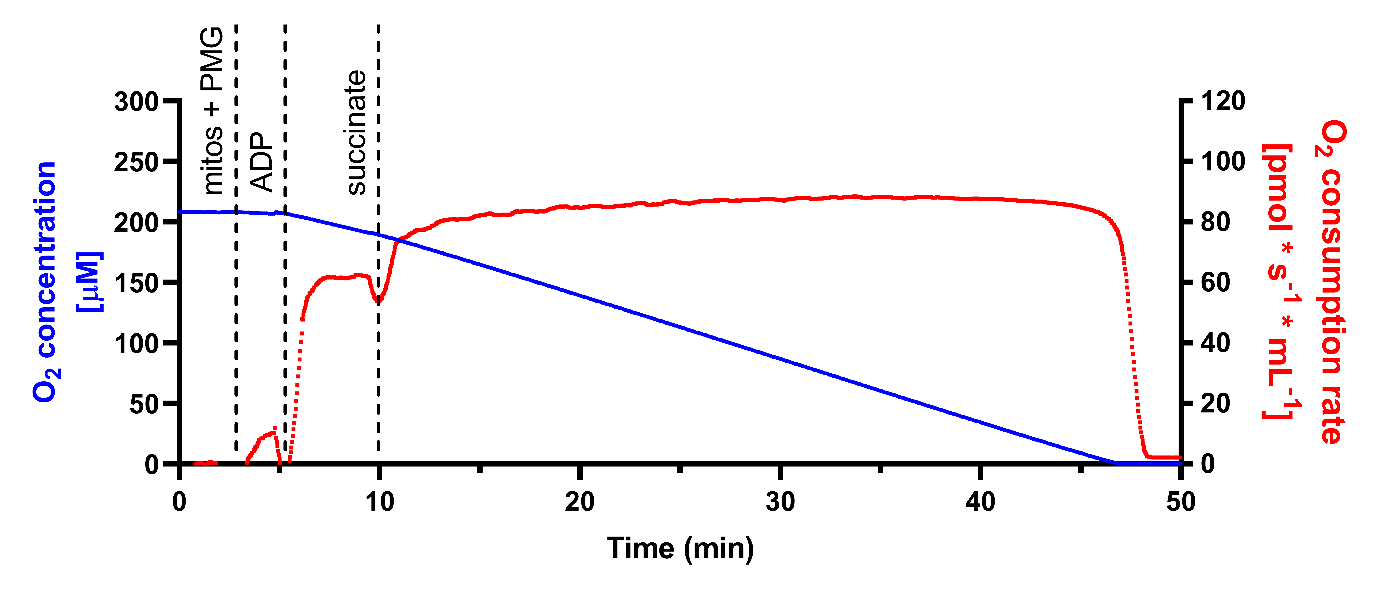

**Supplementary Figure 6. Oxygen consumption recorded in carotid body mitochondrial (mitos) sample when the chamber was fully oxygenated at the start of the protocol.**
Carotid body mitochondrial oxygen consumption remains stable until oxygen concentration reaches very low levels. ADP - adenosine 5′-diphosphate, PMG – pyruvate, malate, glutamate.

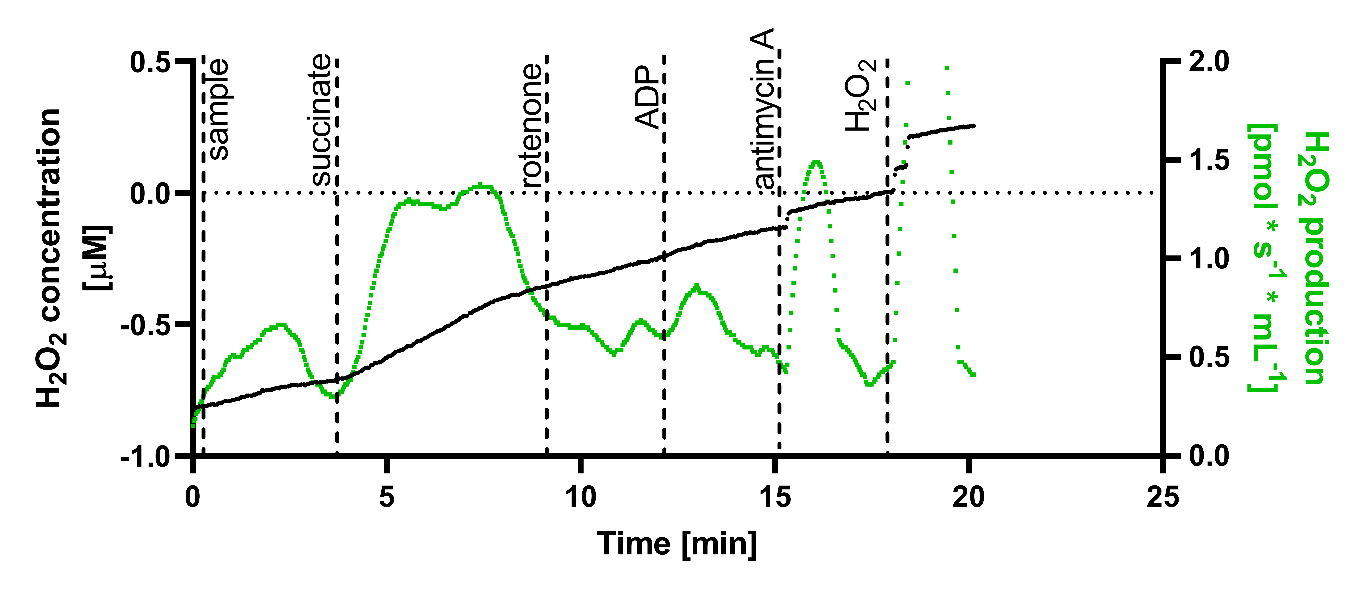

Supplementary Figure 7. Representative trace of the reverse electron transport protocol showing changes in H_2_O_2_ production rate.

ADP - adenosine 5′-diphosphate.

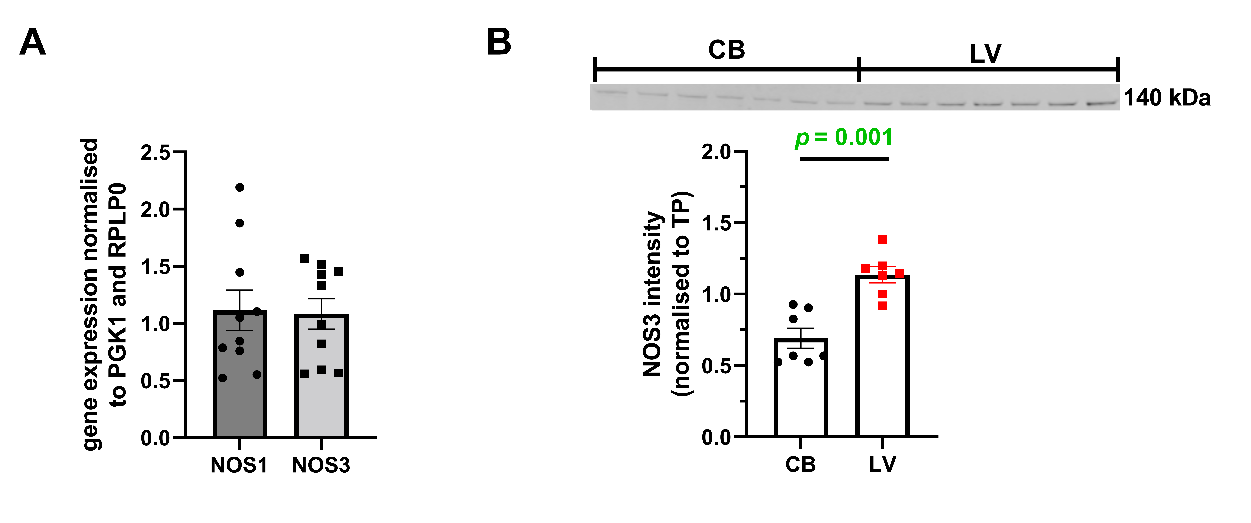

Supplementary Figure 8. Gene expression of neuronal (NOS1) and endothelial nitric oxide synthase (NOS3) in the carotid body (CB; A) and protein abundance of NOS3 (B) in CB and left ventricle (LV) tissue.
A. n = 10 in each group; B. n = 7 in each group, average of 3 technical repeats, parametric unpaired t-test (B).

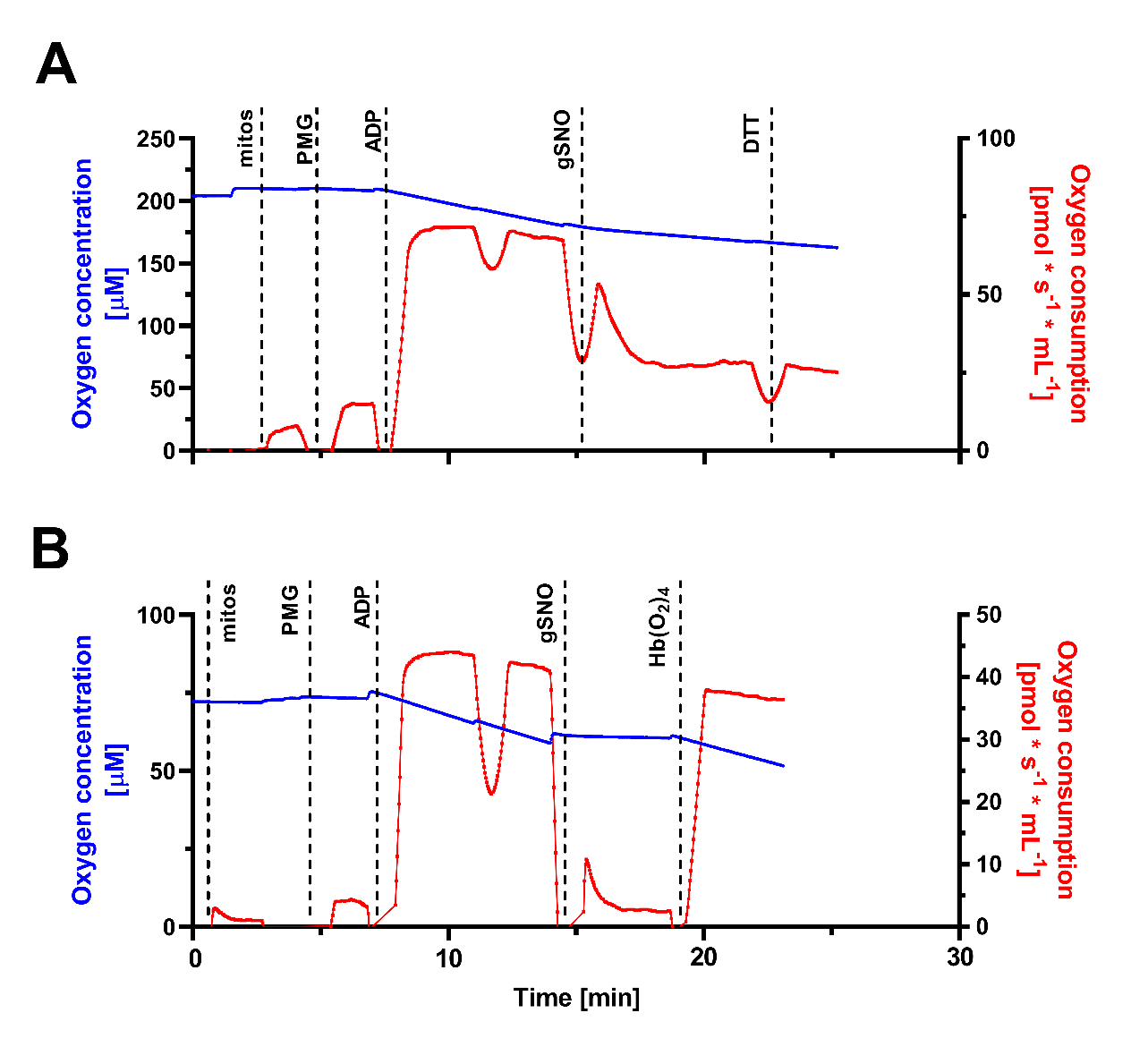

Supplementary Figure 9. Representative traces of protocol 1 (A) and protocol 2 (B).

ADP - adenosine 5′-diphosphate, DTT - DL-Dithiothreitol, GSNO - S-Nitrosoglutathione, Hb(O_2_)_4_ – oxyheamoglobin, mitos – mitochondrial samples, PMG –pyruvate, malate, glutamate.

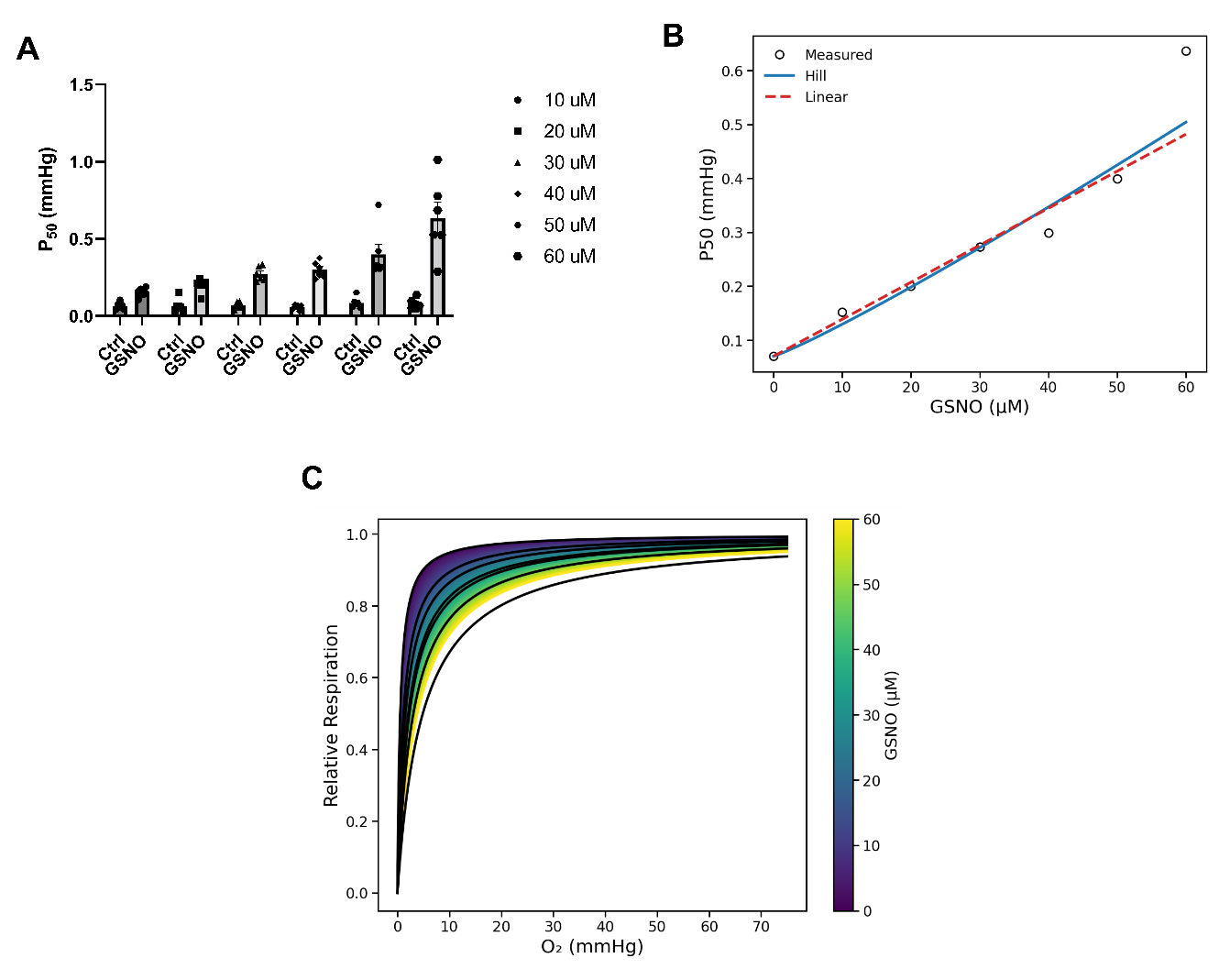

Supplementary Figure 10. GSNO can modulate oxygen affinity of the carotid body mitochondria in a concentration-dependent manner.
A. the effect of varying concentrations (10-60 µM) of GSNO on carotid body complex IV oxygen affinity (P_50_); B. the relationship between the P_50_ value and GSNO concentration; C. modelled relationship between relative respiration and oxygen tension based on the effects of varying concentrations of GSNO on isolated mitochondria.

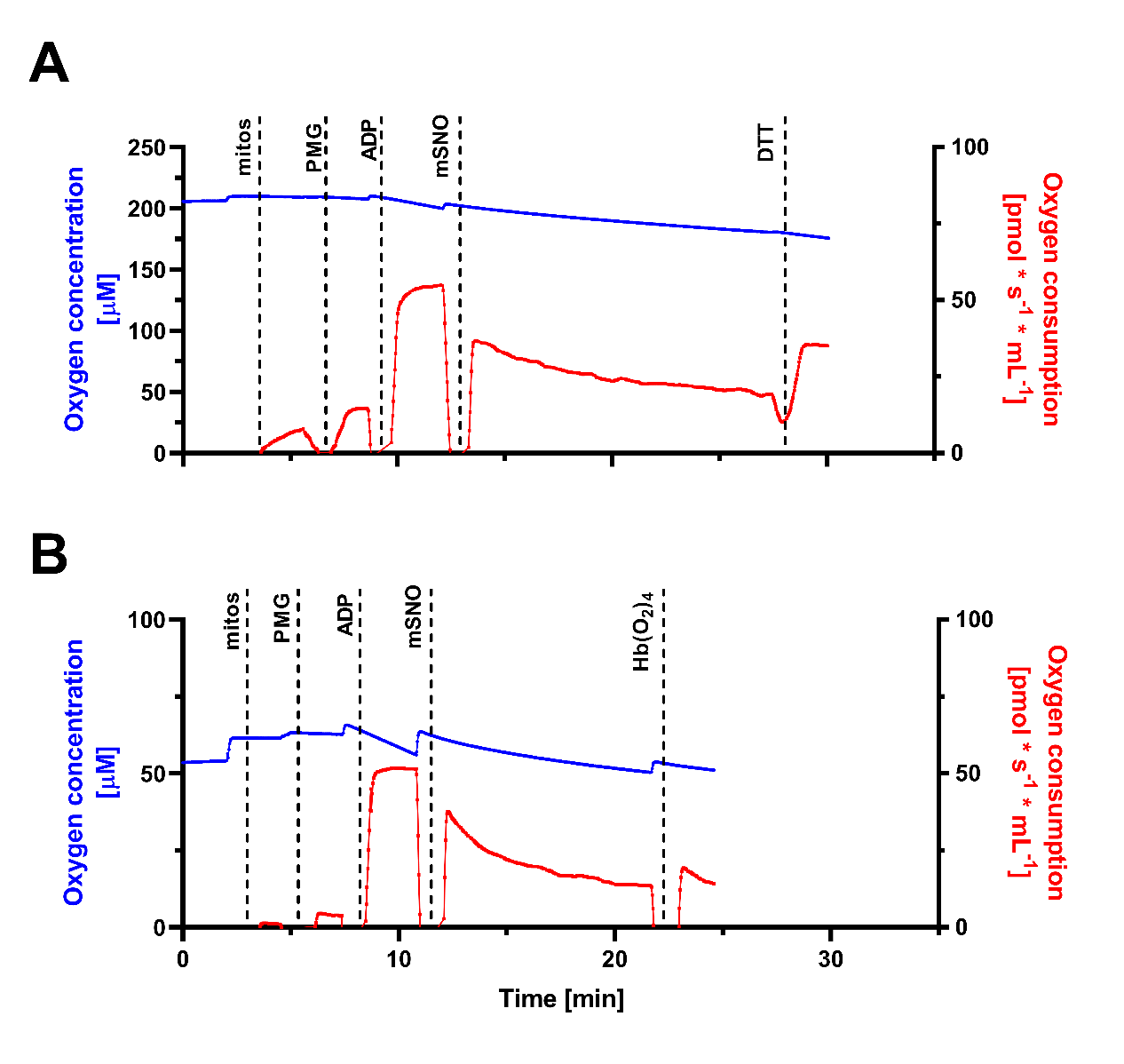

Supplementary Figure 11. Example traces of protocol 1 (A) and protocol 2 (B) with the use of mSNO.

ADP - adenosine 5′-diphosphate, DTT - DL-Dithiothreitol, mSNO - mitoSNO, Hb(O_2_)_4_ – oxyheamoglobin, mitos – mitochondrial samples, PMG –pyruvate, malate, glutamate.

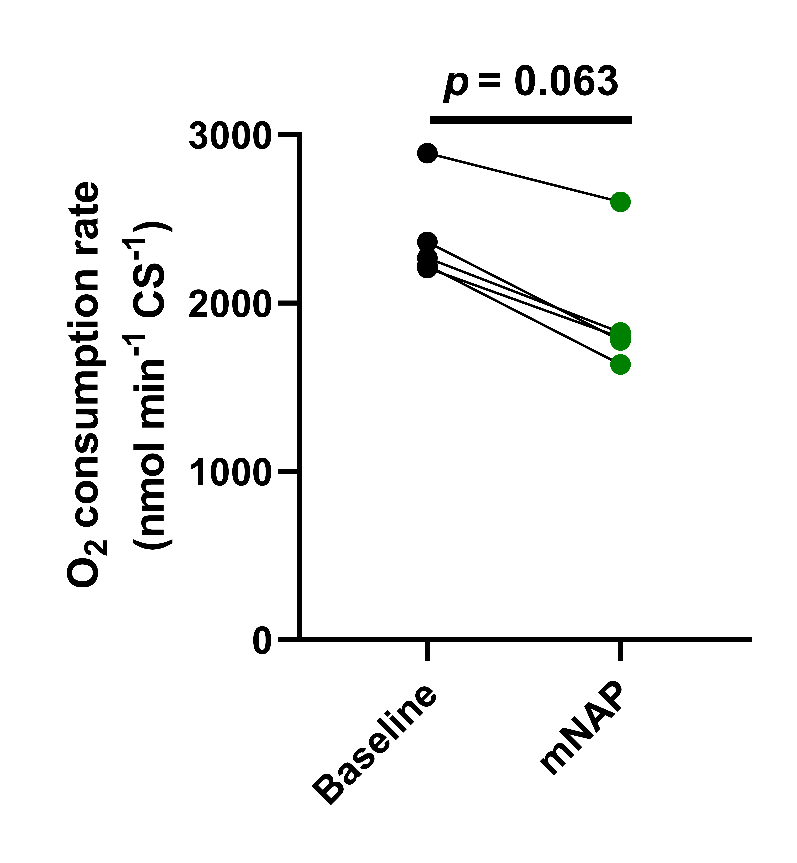

Supplementary Figure 12. The effects of mNAP on respiration in the carotid body mitochondria.
n= 5 in each group, nonparametric paired t-test.

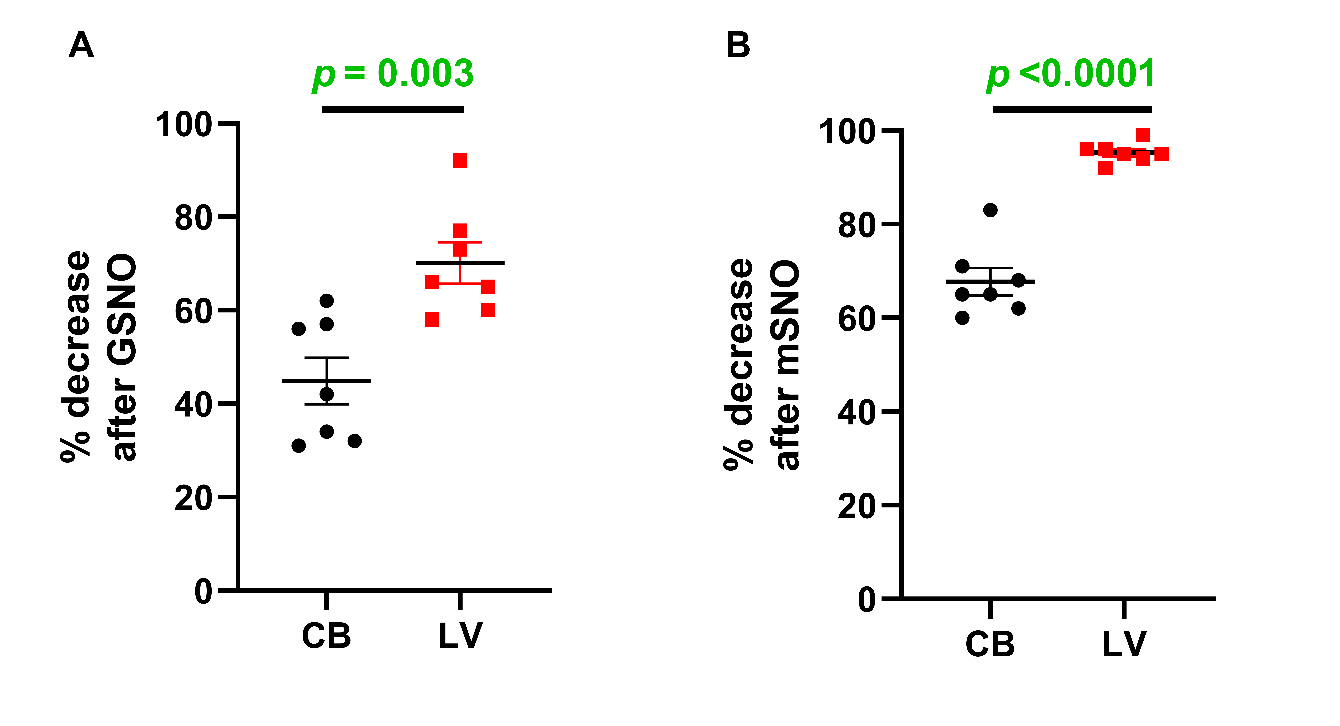

Supplementary Figure 13. The magnitude of GSNO and mSNO inhibition of mitochondrial respiration in carotid body (CB; A) and left ventricle (LV; B).
n = 7 in each group, parametric paired t-test.

1.	Smith CER, Quinn CJ, Clarke JD, Sultan Z, Najem H, Denham NC, Hutchings DC, Whitley AS, Madders GWP, Caldwell JL, et al. Atrial t-tubules adopt a distinct developmental state as Ca^2+^ handling matures postnatally. *J Mol Cell Cardiol*. 2026;212:60-74. doi: 10.1016/j.yjmcc.2025.12.012
